# Long-term dietary interventions fail to mitigate functional connectivity loss and cognitive decline in the TgF344-AD rat model of Alzheimer’s Disease

**DOI:** 10.1101/2025.08.25.672063

**Authors:** Judith R.A. van Rooij, Monica van den Berg, Manoy Van Vosselen, Elke Calus, Tamara Vasilkovska, Lauren Kosten, Ignace Van Spilbeeck, Johan Van Audekerke, Debby Van Dam, Daniele Bertoglio, Mohit H. Adhikari, Marleen Verhoye

**Affiliations:** Bio-Imaging lab, Faculty of Pharmaceutical, Biomedical and Veterinary Sciences, University of Antwerp, Antwerp, Belgium; µNEURO Research Centre of Excellence, University of Antwerp, Antwerp, Belgium; Laboratory of Neurochemistry and Behavior, Department of Biomedical Sciences, University of Antwerp, Antwerp, Belgium; Department of Neurology and Alzheimer Center, University Medical Center Groningen, University of Groningen, Groningen, The Netherlands

**Keywords:** caloric restriction, resveratrol, Alzheimer’s Disease, resting-state functional MRI, cognitive dysfunction, amyloid

## Abstract

**Introduction:** Alzheimer’s disease (AD) is a complex neurodegenerative disorder, characterized by altered brain functional connectivity and network activity, detectable using resting-state functional MRI (rsfMRI). Caloric restriction (CR) and its mimetic resveratrol (Rsv) have shown potential in reducing AD-related pathology and preserving brain function, though research on long-term efficacy is limited. Therefore, we aimed to assess the effects of short-term CR and Rsv administrations on resting-state (rs) functional connectivity (FC), after which we assessed the effects of long-term CR and Rsv interventions on rs-FC, spatial memory, amyloid burden, and neuroinflammation in TgF344-AD rats (Tg) and wild-type (WT) littermates.

**Materials and Methods:** We used rsfMRI to investigate rs-FC changes after CR (40%) or daily Rsv supplementation (10 mg/kg, oral) in male and female, WT and TgF344-AD (Tg) rats at the age of 4 months and 11 months, after, respectively, 1 month (short-term) and 8 months (long-term) of either Rsv supplementation, control (Ctrl) or CR diet. Additionally, spatial memory was assessed utilizing the Morris Water Maze (MWM), followed by histological validation of amyloid plaque load (X34), astrogliosis (GFAP) and microgliosis (IBA-1) at 11 months of age.

**Results:** Both long-term CR or Rsv supplementation exert limited effects in the TgF344-AD model on the known AD-related pathological hallmarks. Short-term CR led to a reduction in rs-FC in female TgF344-AD rats compared to Rsv Tg supplemented rats for connections primarily within and between the hippocampal network, while between other connections, CR reduced rs-FC compared to Ctrl in females, irrespective of genotype. Rsv increased rs-FC compared to CR for only a few connections in females, again irrespective of genotype. Long-term CR decreased rs-FC in male Tg CR rats, compared to Tg Ctrl and WT CR rats, primarily for connections within the lateral cortical network (LCN). For other connections, CR reduced rs- FC when compared to Ctrl, irrespective of genotype. Overall, Rsv supplementation showed negligible effects on rs-FC. Moreover, long-term CR yielded modest cognitive improvements in male Tg rats, as evidenced by enhanced performance in the MWM but not in females. Histological validation after long-term dietary intervention revealed region-specific increase or decrease of amyloid burden after CR or Rsv supplementation, respectively. Additionally, CR reduced IBA-1 levels in males, irrespective of genotype, GFAP levels were unaffected by long-term dietary intervention.

**Conclusion:** Altogether, our findings indicate that long-term CR and Rsv exert distinct and limited effects on AD-related pathology in the TgF344-AD model, with CR demonstrating a modest but greater effect on functional and cognitive measures compared to Rsv. This study underscores the difficulty of altering key disease processes with these long-term dietary approaches, highlighting the need for more comprehensive long-term studies to elucidate the potential modulatory role of dietary interventions on AD pathophysiology.

## 1. Introduction

Alzheimer’s disease (AD), the most common cause of dementia, is one of the most impactful brain disorders in elderly humans, affecting millions worldwide. AD is characterized by gradually progressive neurological and psychological deficits caused by cortical and hippocampal atrophy (1), and impaired brain activity as a result of synaptic loss (2). AD is hallmarked by accumulation of amyloid-beta (Aβ), the formation of tau neurofibrillary tangles (NFTs), and neuroinflammation, which all collectively contribute to cognitive decline (3).

Functional brain changes related to AD pathology are present across the clinical spectrum, where especially important is the presence of early perturbations of network activity, prior to the start of amyloid deposition (4). These changes have been uncovered through utilization of resting-state functional MRI (rsfMRI) (5). RsfMRI is a non-invasive neuroimaging modality that measures low-frequency fluctuations in brain activity in the absence of explicit tasks or stimuli by detecting blood oxygenation level-dependent (BOLD) signal fluctuations. Specific spatially distributed brain regions, forming the resting-state networks (RSNs), present a high temporal correlation between their BOLD signal, defined as functional connectivity (FC) (6–8).

Clinical rsfMRI studies in patients with (prodromal) AD have shown FC alterations, primarily within RSNs such as the default mode network (DMN), typically anticorrelated to the frontoparietal network (FPN) (9–13). In mouse models of amyloidosis and tauopathy, we and others have observed early hyperconnectivity within- and between regions of the hippocampus and the rodent analogue of the DMN (the default-mode-like network (DMLN)) in pre- and early-plaque stages, followed by hypoconnectivity once plaque deposition was prominent (14–18). In the TgF344-AD rat model of AD, characterized by the presence of human APP_swe_ and PS1_ΔE9_ mutations, we and others have observed aberrant DMLN connectivity patterns at pre- and early-plaque stages (19, 20) linked to cognitive and behavioral performance (21–23), highlighting the promise of rs-FC as sensitive early indicators of AD-related neurofunctional alterations.

Despite decades of preclinical research and numerous clinical trials, effective treatment of AD is still lacking, mostly due to incomplete insights into the disease mechanisms and late-stage diagnosis of AD (5). Caloric restriction (CR), defined as a reduction in energy intake without malnutrition, has demonstrated significant benefits in clinical AD studies (24). In rodent models of AD, short- (6-16 weeks) and long-term (8-15 months) CR has shown to reduce activation of astrocytes (25), as well as improved spatial memory (26), reduced Aβ plaque deposition (27–29), attenuation of astrogliosis and reduction of tau hyperphosphorylation (30). However, long-term adherence to CR poses a major challenge, limiting its practical application in clinical settings. To address this limitation, CR-mimetics such as resveratrol (Rsv) have been introduced. Rsv is a polyphenolic compound naturally found in foods like berries and peanuts. In rodent models of AD, Rsv has been linked to reductions of amyloid burden (31, 32) through direct disruption of Aβ aggregation (33, 34), protection of blood-brain barrier integrity (32) and prevention of memory loss after long-term Rsv supplementation (35). Altogether, CR and Rsv offer promising avenues for preserving brain function and reducing the risk or progression of neurodegeneration in AD.

CR or Rsv-induced changes in FC have been investigated using rsfMRI. In healthy older adults, short-term CR (12 weeks) has shown to reduce FC between regions of the hippocampal network (Hipp) and the FPN (36), while on the contrary, short-term Rsv supplementation (26 weeks) increased FC between regions of the Hipp, the DMN, and the FPN (37, 38). In our previous work, short-term CR and Rsv supplementation (4 weeks) has shown to induce a female-specific decrease in healthy young adult rats of FC between the Hipp and the subcortical network (SubC) and the DMLN (39). Moreover, patients with mild cognitive impairment (MCI) who had received short-term Rsv supplementation (26 weeks), showed increased FC between the hippocampus and angular cortex (40).

Despite these findings, there is a lack of knowledge regarding the potential benefits of long-term CR or Rsv, especially in the context of neurodegenerative diseases such as AD. To address these shortcomings, we first aimed to assess and compare effects of short-term (1 month long) CR and Rsv interventions on rs-FC alterations in 4-month (4M) old TgF344-AD rats and WT littermates. Then, we aimed to investigate the effects of long-term (8-months long) interventions on alterations in rs-FC and spatial memory and sought histological validation of their effects on amyloid burden and neuroinflammation in 11 month-old (11M) TgF344-AD rats and WT littermates.

## 2. Materials and methods

### 2.1. Animals

Part of the data utilized in this study (the 4M rsfMRI data acquired in WT rats) were initially obtained and are published in a previous manuscript (39). All F344 (WT) (RRID: RGD_60994, Charles River, Italy) and TgF344-AD (Tg) rats (RRID:RGD_10401208, RRRC) used within this study were bred in-house. All rats were kept under controlled environmental conditions (12-h light/dark cycle, (22 ± 2)°C, 40-60% humidity). Water was provided ad libitum. All procedures were in accordance with the European Community Council Directive on the protection of animals used for scientific purposes (2010/63/EU), and were approved by the Committee on Animal Care and Use at the University of Antwerp, Belgium (ECD: 2021-59).

### 2.2. Dietary treatment paradigm

A total of 84 male and female WT and Tg rats (n = 7/sex, genotype, treatment) were randomly assigned to one of three groups (Figure 1) receiving 40% CR, Rsv supplementation, or serving as controls (Ctrl).

**Figure 1:**
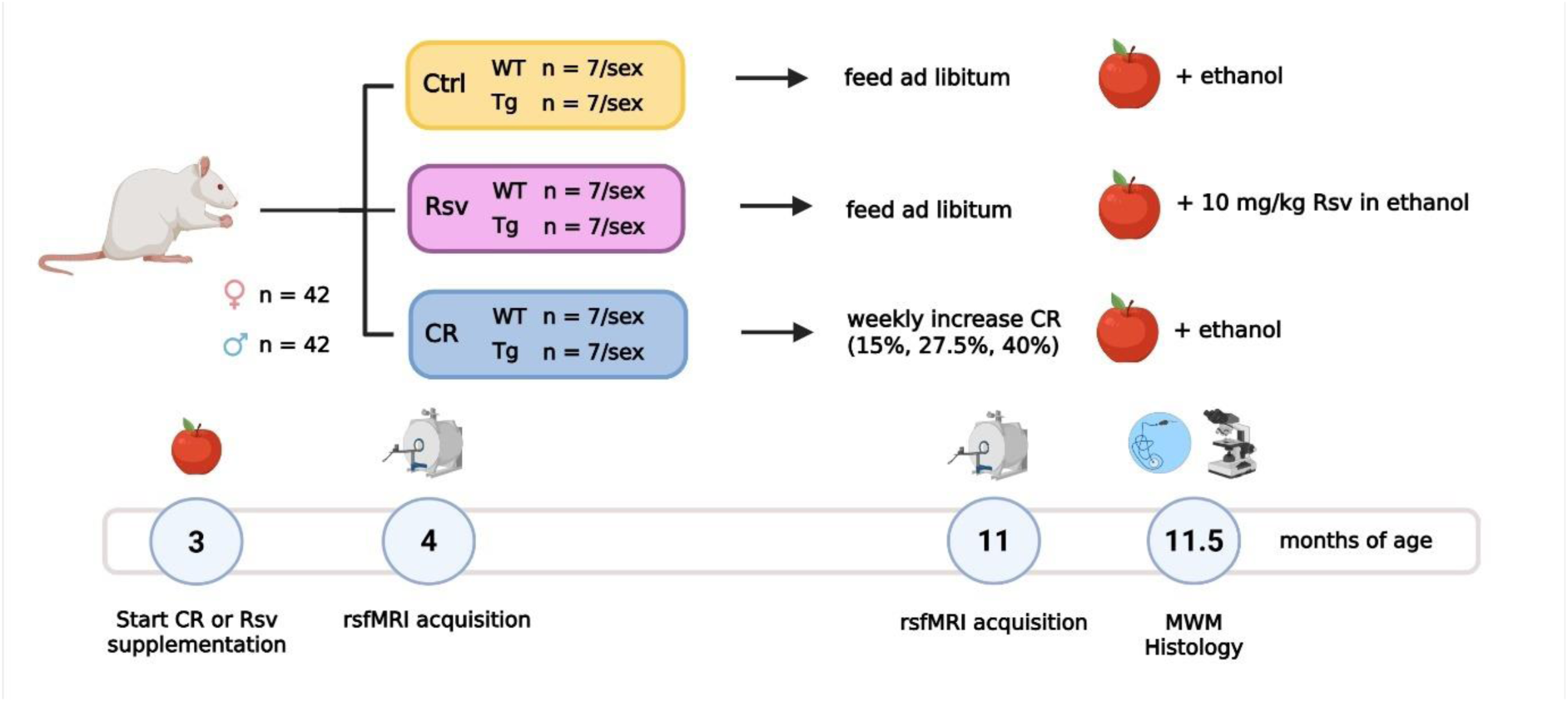
Overview of the experimental dietary treatment paradigm and study design. Number of animals per sex, genotype and group are given. Dietary intervention and supplementation of the apple chips starts at three months of age (3M). RsfMRI data are acquired at the (4M) and eleven (11M) months of age. After the 11M rsfMRI data acquisition, spatial memory was assessed using the MWM. Afterwards, animals were sacrificed at 11.5 months of age for ex vivo analysis. The apple represents the supplementation of Rsv or its vehicle using dried apple chips. Created with Biorender.com

All rats were single housed and received treatment (CR, Rsv supplementation, or vehicle only supplementation) from three months of age for the duration of eight months, until the age of 11 months. Rsv supplementation (10 mg/kg, oral, High Potency Trans-Resveratrol 600, Doctor’s Best®, California, USA), and CR was implemented as described by van Rooij et al. (39). CR rats were exposed to a gradual weekly decrease in caloric uptake until the decrease reached 40%. To allow adaptation to restricted feeding, the animals were fed one week with 15% CR, a second week with 27.5% CR and 40% CR thereafter. Food fortified with vitamins and minerals was provided to avoid malnutrition (Rat/Mouse Fortified, Ssniff®, Germany). CR of 15%, 27.5% and 40% was calculated based on averaged food intake of Ctrl (male/female n = 3/3) and Rsv (male/female n = 3/2) rats, adjusted for age and sex. Ctrl and Rsv supplemented rats were provided standard chow (Ssniff®, Germany) ad libitum. Rsv solubilized in 99.8% ethanol (stock: 50 mg/ml, Thermo Fisher Scientific), was applied to dried apple chips (JR Farm, Germany) and supplemented daily to the rats in the Rsv group. To minimize the difference between conditions, all CR and Ctrl rats were supplemented with a daily dose of dried apple chips with vehicle only (99.8% ethanol). This volume of ethanol was determined through averaging the total daily Rsv volumes of all Rsv supplemented rats, adjusted for age and sex. Rats were weighed prior to the start of the treatment and each week afterwards for the entire duration of the experiment (Supplementary Figure 1). The percentage body weight (BW) change respective to the starting weight was calculated for each subject.

### 2.3. rsfMRI acquisition

RsfMRI data of all rats were acquired at 4M and 11M, corresponding to, respectively, short-term (1 month long), and long-term (8-month long) CR or Rsv supplementation. Rats were anaesthetized using 5% isoflurane (IsoFlo®, Zoetis, US), administered with a gaseous mixture of 200 ml/min O_2_ and 400 ml/min N_2_. For further animal handling and positioning, the isoflurane level was reduced to 3% after which a subcutaneous bolus injection of medetomidine (0.05 mg/kg, Domitor®, Vetoquinol, France) was administered. Continuous subcutaneous infusion of medetomidine (0.1 mg/kg/h) started 15 minutes after bolus administration, with the isoflurane level being lowered to 0.4%, to be maintained throughout the whole MRI session. Physiological parameters (breathing rate, heart rate, O_2_ saturation and body temperature) of the animal were closely monitored (MR-compatible Small Animal Monitoring, Gating, and heating system, SA Instruments). Body temperature was maintained (37.0 ± 0.5 °C) using a feedback-controlled warm air circuitry (MR-compatible Small Animal Heating System, SA Instruments Inc., USA).

MRI data were acquired on a 7T Pharmascan MR scanner (Bruker®, Germany) with a volume transmit RF coil and a 2x2 array head receiver RF coil. To ensure uniform slice positioning between subjects, multi-slice T2-weighted (T2w) TurboRARE images were acquired in three directions (echo time (TE): 33 ms, repetition time (TR): 1800 ms, RARE factor: 8, field of view (FOV): (35 x 35) mm^2^, matrix: [256 x 256]). Whole-brain rsfMRI data were acquired 40 minutes after the medetomidine bolus by using a single shot gradient echo echo-planar imaging (EPI) sequence (TE: 18 ms, TR: 600 ms, FOV (30 x 30) mm^2^, matrix [96 x 96], 12 coronal slices of 1 mm, slice gap: 0.1 mm, 1000 repetitions), for a total of 10 minutes. An anatomical 3D image was acquired for registration purposes, with a T2w-TurboRARE sequence (TR: 1800 ms, TE: 36 ms, RARE factor: 16, FOV: (35 x 35 x 16) mm^3^, acquisition matrix: [256 x 256 x 32], reconstruction matrix: [256 x 256 x 64]). At the end of the scan session, a subcutaneous injection of 0.1 mg/kg atipamezole (Antisedan®, Pfizer, Germany) was administered to counteract the effects of the medetomidine anesthesia and the animals were allowed to recover under a heating lamp. All animals recovered within 15-20 minutes after the end of the scan session.

### 2.4. rsfMRI image preprocessing

All preprocessing steps were performed with MATLAB R2020a (Mathworks, Natick, MA) and ANTs (Advanced Normalization Tools). The rsfMRI data were padded using an in-house MATLAB script. Debiasing, realignment, co-registration, normalization and smoothing of the data were performed using SPM 12 software (Statistical Parametric Mapping). First, subject-specific 3Ds were debiased after which a study specific 3D-template was created from a subset of animals (n = 48 (n = 3/sex, treatment, age, genotype)) in ANTs. All individual 3Ds were normalized to the study-specific 3D template using a global 12-parameter affine transformation followed by a non-linear transformation. The rsfMRI EPI images were realigned to the first EPI image using a 6-parameter rigid body transformation estimated using a least-squares approach. Data acquired at either 4M or 11M of age were independently assessed and included for further analysis if within the realignment step, image shifts within the image series were less than once the voxel size (= 0.3125 mm). Here, data acquired at 4M for one female WT Ctrl rat was excluded from further analysis, due to image shifts larger than once the voxel size. RsfMRI data were co-registered to the animal’s respective 3D image using a global 12-parameter affine transformation with mutual information used as similarity metric and normalized to the study-specific template using the earlier estimated transformation parameters of the normalization of the individual subjects 3D image to the template. RsfMRI data were smoothed in-plane using a Gaussian kernel with full width at half maximum of twice the voxel size and filtered (0.01 – 0.2 Hz) with a Butterworth band- pass filter, after which quadratic detrending was performed on the filtered images. As a quality check of the data, a subject level independent component analysis (ICA) was performed to assess subject level FC. FC maps (20 components) that showed clear bilateral components were included in further data analysis. No subject data acquired at 4M of 11M was excluded following this quality check.

### 2.5. FC analysis

Region of interest (ROI)-based FC analysis was performed on the preprocessed rsfMRI data. A neuroanatomical atlas comprised of 71 anatomical parcels (Fischer 344, https://www.nearlab.xyz/fischer344atlas), was warped onto the study-specific template and down- sampled (ANTs) to match the EPI space dimensions. Out of the 71 parcels, 47 unilateral grey matter ROIs (for both the left (L) and right (R) hemisphere for each region) were selected, excluding regions sensitive to susceptibility artefacts, partial volume effects and small size (Supplementary Table 1). The selected ROIs represent five prominent rodent RSNs: the default mode like network (DMLN), the hippocampal network (Hipp), the sensory network (Sens), the lateral cortical network (LCN) and the subcortical network (SubC) (Supplementary Table 1). For each subject, Pearson correlation coefficients between the ROI-averaged BOLD signal timeseries of each pair of ROIs were calculated and Fisher z-transformed yielding subject-wise [47x47] FC matrices. Afterwards, the subject-wise matrices were averaged per age, sex, genotype and treatment, yielding group-wise [47x47] FC matrices (Supplementary Figures 2 and 3).

### 2.6. Hidden-platform Morris Water Maze test

Spatial learning was assessed at 11M using the Morris Water Maze (MWM). The MWM consisted of a circular pool (diameter: 150 cm, height: 30 cm) with a blacked-out bottom, filled with water kept at 25°C. The pool was divided into four quadrants: NE, SW, SE, NW. A round, acrylic glass platform (diameter: 15 cm) was placed in a fixed position at the center of the NE quadrant and hidden 1 cm beneath the water surface. The set-up was surrounded by white walls with high contrast 2D visual cues surrounding the maze. MWM training and acquisition was performed over a period of 4 days and consisted of two daily trial blocks (one at 9.00 am and one at 13.00 pm) consisting of the 4 trials each with a 15-minute inter-trial interval. The starting positions were based on the quadrants and were consistent across all animals, while varying in a randomized order per trial. Animals were placed in the pool with their nose towards the wall. In case an animal was unable to reach the platform within 120 seconds, it was guided towards the platform where it had to remain for 15 seconds before being returned to its home cage. For each trial, path length, escape latency and swim speed were determined, and afterwards the total path length, escape latency and average velocity was calculated encompassing all trials (41). During the probe trial, performed three days after the final acquisition trial block, the platform was removed from the maze and each animal was allowed to swim freely for 100 seconds. Afterwards, we determined the swim speed, path length, the percentage of time spent in the quadrant of the pool that contained the hidden platform and the number of times the rats crossed the former platform’s coordinates. Animals’ trajectories during the training days and probe trail were recorded using a computerized video-tracking system (Ethovision 14, Noldus, Wageningen, the Netherlands). Four subjects (1 WT Ctrl male, 1 WT CR male, 1 Tg Ctrl male, 1 Tg Ctrl female) were excluded due to visual impairment or high levels of anxiety limiting their ability to perform the MWM.

### 2.7. Immunohistochemical staining, acquisition and analysis

To evaluate the effects of dietary intervention on amyloid accumulation and neuroinflammation histological analyses were performed on cryosections of 36 TgF344-AD rats (Ctrl n = 6/sex, Rsv n = 6/sex, CR n = 6/sex) and 24 WT littermates (Ctrl n=4/sex, Rsv n = 4/sex, CR n = 4/sex). Rats were deeply anaesthetized using an intravenous injection of pentobarbital (Dolethal®, Vetoquinol, Belgium). Cardiac perfusion was performed with an ice-cold PBS solution. Brains were surgically removed and hemispheres were separated. All right hemispheres were post-fixated in a 4% paraformaldehyde solution, followed by a sucrose gradient (5%, 10% and 20%). All left hemispheres were immediately snap frozen in liquid nitrogen and stored at −80 °C until further processing upon extraction, while the right hemispheres were snap frozen after completion of the sucrose gradient. For immunohistochemistry, right hemispheres were embedded in an OCT-embedding medium (VWR) for sectioning. At Bregma levels 1.13 and 3.90 mm, sagittal sections of 10 µm were cut using a Leica CM1950 cryomicrotome (Leica BioSystems, Belgium) and thaw-mounted on Epredia Superfrost Plus Adhesion Microscope Slides (Epredia, Breda, Netherlands).

Histological analysis focused on the investigation of the effects of dietary intervention on amyloid burden and neuroinflammation. Aβ plaques were visualized using X-34 (Merck Life Science, Hoeilaart, Belgium). Sections were permeabilized for 15 min with 0.2% Triton in 0.01 M PBS (pH 7.4) prior to a 20-min incubation with x-34 (10 µM) in 40% ethanol (pH 10). Propidium iodide (5 µg/mL; 5 min; Sigma-Aldrich) was used as a nuclear stain. Sections were mounted using a permanent mounting medium Citifluor PVP-1 + Antifadent. Glial cells were visualized using markers for GFAP (staining for reactive astrocytes) and IBA-1 (staining for microglia). Sections were washed and incubated in citrate buffer (Ph 6.0) to unmask antigen, followed by a preincubation with H_2_O_2_ for 5 minutes. Next, sections were incubated for 60 minutes using a blocking buffer containing 10% NGS and 0.2% Triton X-100 before an overnight incubation with the primary antibodies (GFAP (1:1000, Rabbit, Dako, Z033429, RRID: AB_10013382), IBA-1 (1:500, Rabbit, Wako Chemicals, 019-19741, RRID: AB_839504)). After overnight incubation, both sections stained for GFAP or Iba1 were incubated for 60 minutes with a secondary antibody (1:500, Goat, Jackson Immunoresearch, 111-035-006, AB_2337936). Samples were mounted with DPX (Sigma-Aldrich, Germany).

Images of the immunohistochemical (GFAP, IBA-1) and -fluorescent (X34) stained sections were acquired on an Olympus BX51 microscope (Olympus Europa, Hamburg) using a 40x Plan C objective and a DP71 camera. Bright-field images were acquired for GFAP and IBA-1. A X-Cite exacte® light source (Excelitas, United States), a filter set suitable for the applied fluorescent dyes (TxRED (595-615 nm) and BFP (440-447 nm)) was used to visualize X34 and the nuclear counter staining. The software cellSens dimension (version 1.6, Olympus) was used to control image acquisition. Per animal, one image per ROI was acquired on 3 consecutive (on the same glass slide) sagittal sections. Image analysis for was done using QuPath (v0.5.1). ROIs used to calculate the percentage positive staining were manually delineated, excluding image artefacts. An intensity threshold was manually defined per subject and region to measure the positive area percentage of GFAP, IBA-1 and X34 with respect to the delineated region. Positive percentage area X34 for the male and female WT Ctrl rats was near zero and were therefore excluded from further statistical analysis.

### 2.8. Statistics

All statistics were performed per age (4M and 11M) and per sex (male and female) using JMP Pro 17 (SAS Institute Inc.), GraphPad Prism (version 9.4.1. for Windows, GraphPad Software, San Diego, California USA) and MATLAB R2020a (Mathworks, Natick, MA). All data was tested for normality using the Shapiro-Wilk test determining the goodness of fit. All data was normally distributed.

ROI-based FC measures, body weight, percentage positive area GFAP and IBA-1, and behavioral outcomes of the training trials (averaged swim speed, distance traveled and escape latency) and probe trial (number of times platform crossed, time spent in the zone of the platform, swim speed and distance traveled), were subjected to a two-way ANOVA (genotype, treatment, genotype*treatment, FDR corrected p<0.05). Prior to further statistical testing, all FC ROI-pairs that demonstrated either a significant interaction or main effects, were subjected to post-hoc analyses using FDR correction (Benjamini-Hochberg procedure, p<0.05) per observed effect. In case of a significant genotype*treatment interaction, post-hoc tests were performed using Student’s t-test with FDR correction (Benjamini-Hochberg procedure, p<0.05) considering only hypothesis driven relevant comparisons. In the absence of a genotype*treatment interaction, the interaction was removed, and the model was recalculated using only the main effects (genotype and treatment). In case of a significant treatment effect, the three post-hoc comparisons were performed using FDR correction (Benjamini-Hochberg procedure, p<0.05) for the ROI-based FC measures, and a Tukey’s Honestly Significant Difference (HSD) test for the body weight, GFAP, IBA-1 and behavioral outcomes.

Percentage positive area X34 was subjected to a one-way ANOVA (treatment) per region was performed to assess the treatment effects in the TgF344-AD rats. In case of a significant treatment effect, post-hoc tests were performed on all groups using Tukey’s HSD test.

Final graphical representation of the data was created using MATLAB R2020a (Mathworks, Natick, MA), GraphPad Prism (version 9.4.1. for Windows, GraphPad Software, San Diego, California USA), ImageJ (42) and Adobe Illustrator (Adobe Inc.).

## 3. Results

### Short-term dietary intervention with CR lowers ROI-based FC in male and female TgF344-AD rats, but not in WT

First, we aimed to investigate the effects of short-term (1 month) dietary intervention with CR or Rsv supplementation on brain function in 4M old male and female, WT and TgF344-AD rats. Statistical analysis revealed significant (two-way ANOVA, p < 0.05) genotype*treatment interaction effects on ROI-based FC in 102 connections for males and 167 connections for females (Figure 2A, lower diagonal). Only the Tg groups demonstrated a significant between-treatment difference (Figure 2A, upper diagonal) after post-hoc comparisons. A significantly lower FC was observed in male Tg CR rats compared to male Tg Ctrl rats (Figure 2A, left panel, and 2B) between S1 R and FrA R, two regions belonging to the LCN. In female Tg rats (Figure 2A, right panel, and 2C), lower FC was observed in CR rats compared to Rsv supplemented rats, in 41/167 ROI-pairs of which at least one of the regions belonged to the Hipp network.

**Figure 2:**
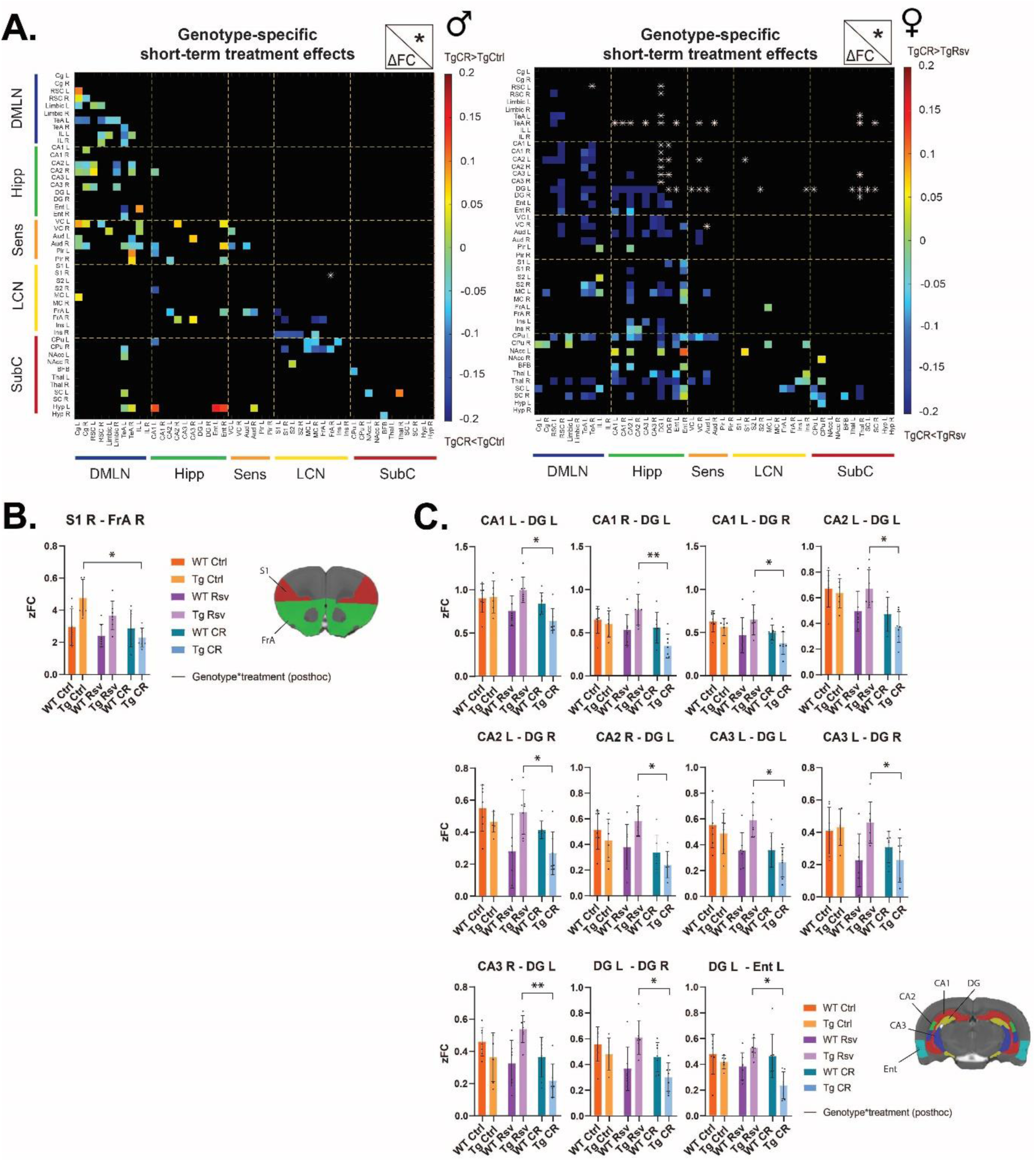
Genotype*treatment interaction effects of short-term (1 month) Rsv supplementation or CR in 4M- old male and female, WT and TgF344-AD rats. A. FC difference (below the diagonal, in color) between male Tg CR and Tg Ctrl rats (A) and between female Tg Rsv and Tg Ctrl rats (B), for connections that showed a significant genotype*treatment interaction effect (p<0.05, two-way ANOVA). Connections with significant inter-treatment FC difference (p < 0.05, post-hoc two-sample T-test, FDR corrected) are indicated above the diagonal by asterisks. B-C. Mean FC (± SD) for all six groups for the connections demonstrating a genotypic specific difference in FC between treatments in 4M males (C) and females (D). Asterisks indicate the levels of statistical significance: *p<0.05, **p<0.01.

Furthermore, we observed main treatment effects (Figure 3A-C) in several connections in both males and females (26 connections for males, 225 connections for females). Post-hoc comparisons, revealed significantly lower FC in female CR rats compared to Ctrl rats for 95 out of 225 ROI-pairs primarily within the DMLN, the LCN, and between the LCN and DMLN, the DMLN and Hipp and between the LCN and SubC (Figure 3B, upper diagonal). These differences were not observed when Rsv supplemented rats were compared to Ctrl (Figure 3A, upper diagonal). Emphasizing the difference between interventions, we observed lower FC in female CR rats when compared to female Rsv rats for 5 out of 225 ROI-pairs between the Hipp and LCN and between Sens and LCN (Figure 3C, upper diagonal). Out of the 26 connections with main treatment effects in 4M old males, none survived post-hoc comparisons with correction. Genotypical differences were observed for 60 connections in 4M old male rats and 60 connections in 4M old female rats (Figure 3D). In males we observed increased FC in Tg rats compared to WT for 52/60 connections within the LCN, and between the DMLN and other RSNs. In females on the contrary, we observed lower FC in Tg rats compared to WT, for 46/60 connections mainly between Sens, DLMN and Hipp regions.

**Figure 3:**
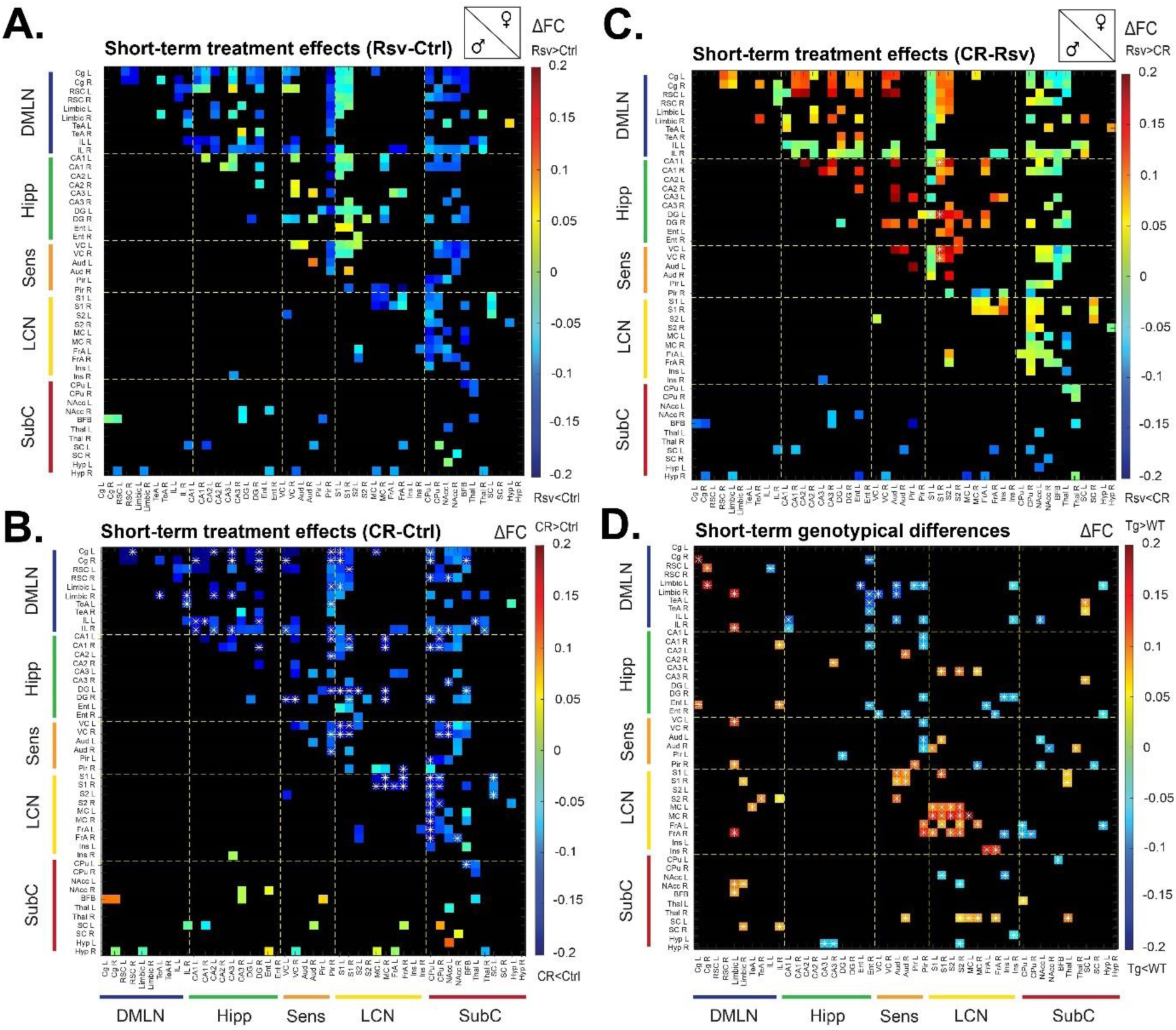
Main effects of short-term (1 month) treatments (Ctrl vs Rsv vs CR) and of genotype (WT vs Tg) on ROI-based FC of 4M old male and female rats. FC difference (in color) for connections with significant main effect of treatment between Ctrl and Rsv (A), Ctrl and CR (B), and Rsv and CR (C), and genotype (D) (p < 0.05, main effects, two-way ANOVA) in male (lower diagonal) and female (upper diagonal) rats. Connections with asterisks indicate a significant inter-treatment difference upon post hoc comparisons (p < 0.05, Tukey HSD for each connection, FDR corrected for all connections with significant treatment effect (A-C)) and a significant main genotype effect upon two-way ANOVA (p < 0.05, FDR corrected for all connections with significant genotype effect (D)).

### Long-term dietary intervention with CR lowers ROI-based FC in male TgF344-AD rats

Next, we aimed to investigate the effects of long-term (8 months) dietary intervention with CR or Rsv supplementation on brain function in 11M old male and female, WT and TgF344-AD rats. Again, several ROI-pairs showed a significant interaction effect (two-way ANOVA, p < 0.05; Figure 4A, lower diagonal) (102 connections for males, 36 connections for females). Post-hoc comparison demonstrated lower FC for 24/102 connections in male Tg CR rats when compared to WT CR for ROI-pairs primarily within the LCN and between the SubC and the DMLN, LCN, Hipp and Sens (Figure 4A, bottom left panel, upper diagonal; Figure 4B). This specific inter-genotypic difference was not observed for Ctrl or Rsv male rats. Moreover, post-hoc comparison demonstrated lower FC in 4/102 connections in male Tg CR rats, compared to Tg Ctrl (Figure 4A, top left panel, upper diagonal; Figure 4B). In females, the only significant post-hoc difference was observed (Figure 4A, top right panel, upper diagonal; Figure 4C) for Pir R – TeA L, showing a significantly lower FC in Tg Ctrl rats compared to WT Ctrl. Interestingly, this difference was not found significant when the Tg rats were compared to WT in either the CR or Rsv groups.

**Figure 4:**
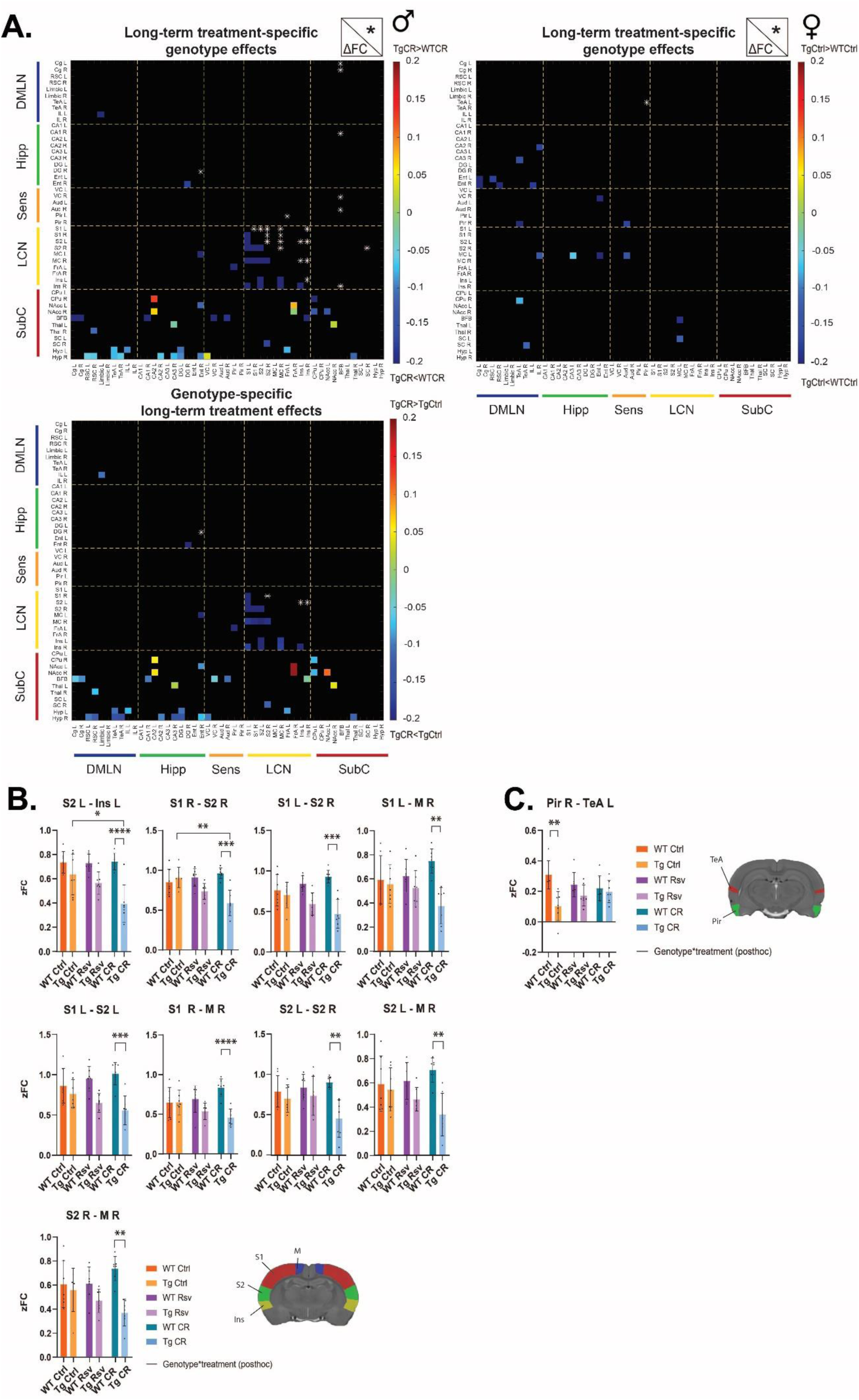
Genotype*treatment interaction effects of long-term (8 months) Rsv supplementation or CR in 11M- old male and female, WT and TgF344-AD rats. FC difference (below the diagonal, in color), between male Tg CR and Tg Ctrl rats (A), male WT and Tg CR rats (B) and female WT and Tg Ctrl rats (C), for connections that showed a significant genotype*treatment interaction effect (p<0.05, two-way ANOVA). In each case, connections with significant FC difference (p < 0.05, post-hoc two-sample T-test, FDR corrected) are indicated above the diagonal by asterisks. B-C. Mean FC (± SD) for all six groups for the connections demonstrating a genotypic specific difference in FC between treatments in 4M males (D) and females (E). Asterisks indicate the levels of statistical significance: *p<0.05, **p<0.01, ***p<0.001, ****p<0.0001.

Moreover, we observed significant main effects of long-term treatments (Figure 5A-C) (178 connections for males, 39 connections for females), irrespective of genotype. Post-hoc comparisons demonstrated lower FC for 45 out of 178 connections in male CR rats when compared to Ctrl for ROI-pairs primarily within- and between the LCN, Sens and DMLN (Figure 5B, lower diagonal). Again, rs-FC differences for these connections were not significant when comparing Rsv supplemented male rats to Ctrl (Figure 5A, lower diagonal), highlighting treatment-specific FC alterations. In females, one singular inter-treatment difference was found significant for the CA3 R – Thal R ROI-pair, revealing higher FC in CR rats compared to Rsv (Figure 5C, upper diagonal). Finally, we observed significant genotypical effects for 440 connections in male and 166 connections in female rats (Figure 5D). In males, we observed decreased FC in Tg rats compared to WT irrespective of treatment, for 432/440 connections. Within and between the SubC and other RSNs we observed an increased FC in Tg rats compared to WT for 8/440 ROI-pairs. In females, we observed lower FC in Tg compared to WT in 165/166 ROI-pairs within- and between all networks.

**Figure 5:**
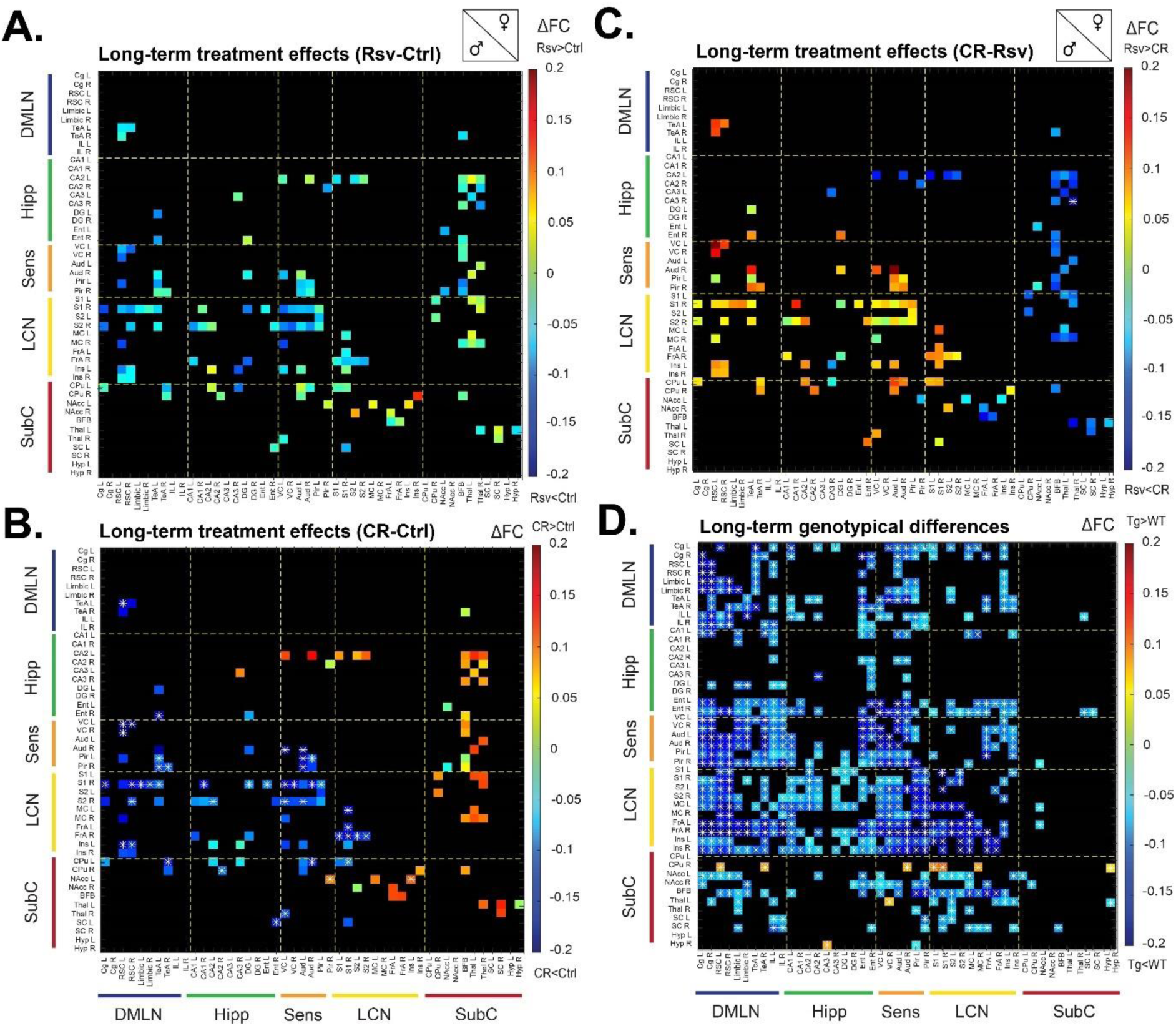
Main effects of long-term (8 months) treatments (Ctrl vs Rsv vs CR) and of genotype (WT vs Tg) on ROI-based FC of 11M old male and female rats. FC difference (in color) for connections with significant main effect of treatment between Ctrl and Rsv (A), Ctrl and CR (B), and Rsv and CR (C), and genotype (D) (p < 0.05, main effects, two-way ANOVA) in male (lower diagonal) and female (upper diagonal) rats. Connections with asterisks indicate a significant inter-treatment difference upon post hoc comparisons (p < 0.05, Tukey HSD for each connection, FDR corrected for all connections with significant treatment effect (A-C)) and a significant main genotype effect upon two-way ANOVA (p < 0.05, FDR corrected for all connections with significant genotype effect (D)).

### Effects of long-term CR or Rsv supplementation on spatial memory in TgF344-AD rats and WT littermates

To assess the potential effects of long-term dietary interventions on cognition in both WT and TgF344- AD rats, spatial memory was assessed using the MWM (Figure 6, Supplementary Figures 4 and 5).

**Figure 6:**
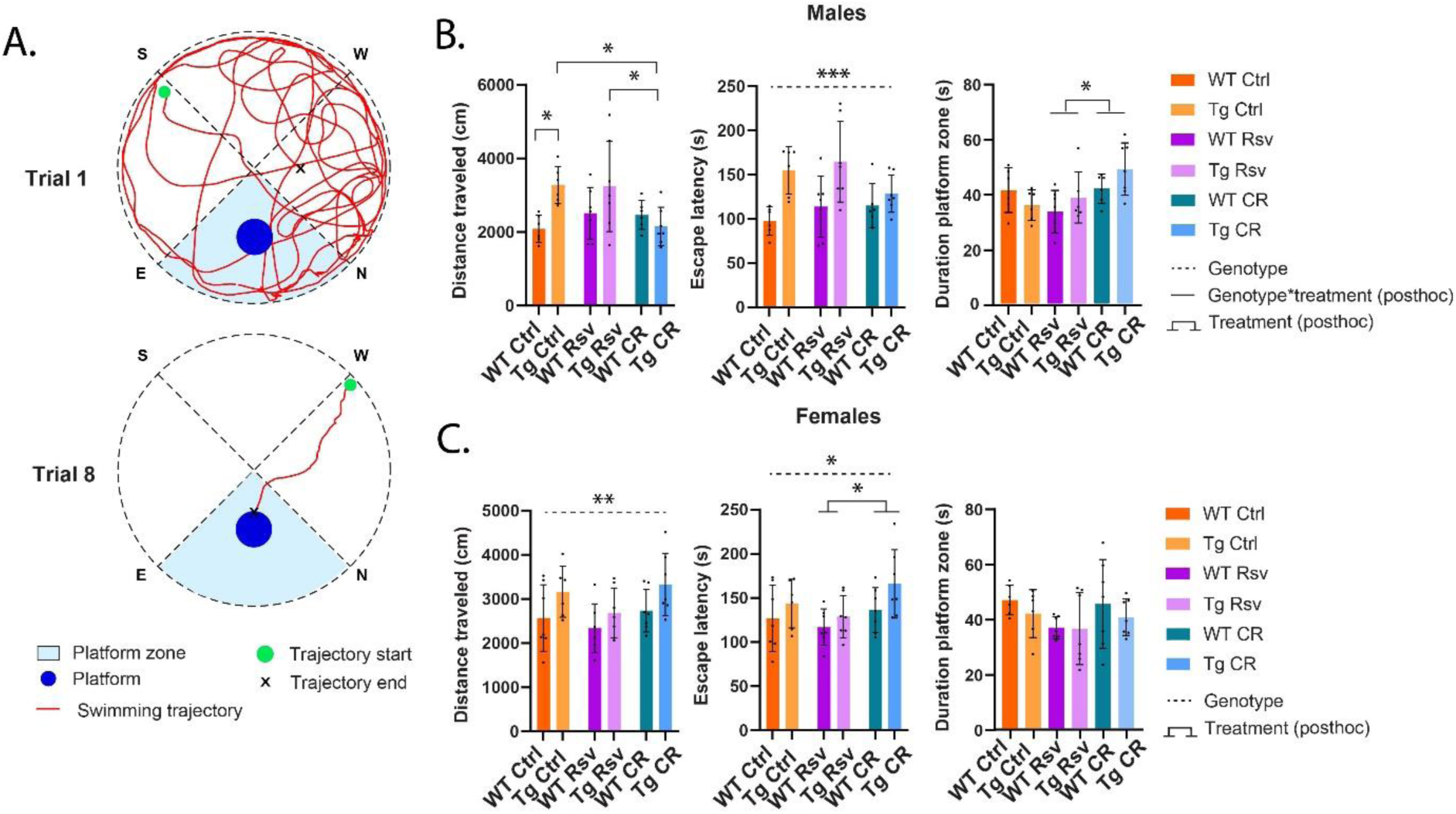
Morris Water Maze (MWM) outcomes reflecting spatial memory. A. Schematic overview of the MWM, divided into four quadrants, with the platform placed between NE. Plotted are example trajectories for trial 1, where longer trajectories are observed, compared to trial 8. Bar plots show the total sum average of the distance traveled, escape latency, during the 8 trials and total duration of time spend in the platform zone during the probe trial for males (B) and females (C). Significant genotype effects are annotated by the black dotted line. Significant treatment effects (post-hoc Tukey HSD p<0.05) are annotated by the black brackets. Significant interaction effects of treatment*genotype (FDR corrected, Benjamin-Hochberg procedure, p<0.05) are noted by the black lines. Asterisks indicate the levels of statistical significance: *p<0.05 **p<0.01, ***p<0.001.

Here, we observed that throughout the learning trials, male Tg Ctrl rats traveled a longer total distance within the maze to reach the platform compared to WT Ctrl rats (p = 0.021)(Figure 6B, left). Interestingly, Tg CR male rats traversed a significantly shorter distance to reach the platform compared to Tg Rsv (p = 0.021) and Tg Ctrl (p = 0.021) rats, which was not observed when compared to WT Ctrl, suggesting a modest improved performance of Tg CR male rats. These observations however were not reflected in the escape latency (Figure 6B, middle), which was significantly longer for Tg male rats compared to WT (p = 0.0003), irrespective of treatment. During the probe trial, male CR rats spent more time in the designated platform zone compared to Rsv rats (p = 0.0143), irrespective of genotype (Figure 6B, right).

In females, Tg rats traveled significantly longer distance (p = 0.01)(Figure 6C, left) and demonstrated significantly longer escape latency (p = 0.0402) compared to WT, irrespective of treatment (Figure 6C, middle). Furthermore, female CR rats took significantly longer to find the platform compared to Rsv supplemented rats (p = 0.0379), irrespective of genotype. No differences were observed in the time spent in the designated platform zone.

### Long-term dietary intervention with CR or Rsv supplementation differentially modulates amyloid-burden in TgF344-AD rats

To investigate the effects of long-term dietary interventions on amyloid burden in TgF344-AD male and female rats, immunofluorescence was performed using X34 to quantify amyloid-plaques (Figure 7A) in 11 different brain regions (Supplementary Table 2) known to be implicated in disease pathology. Positive percentage area X34 for the male and female WT Ctrl rats was near zero (Figure 7A) and were therefore excluded from further statistical analysis. Statistical analysis (one-way ANOVA, treatment, p < 0.05, Tukey HSD, Supplementary Table 3) revealed significant treatment effects in males for 4 out of 11 regions, and 1 out of 11 for females (Figure 7B, Supplementary Figure 6). Post-hoc analysis revealed a decrease of amyloid burden in Tg Rsv supplemented male rats when compared to Tg CR in the CA3 (p = 0.0047) and Tg Ctrl in both the CA3 (p = 0.0079) and Ent (p = 0.0355). In contrast, Tg Rsv male supplemented rats had higher amyloid burden when compared to Tg Ctrl in the MC (p = 0.0153) and Tg CR in the VC (p = 0.0423) and MC (p = 0.0048). Lower accumulation of amyloid was observed in male Tg CR rats when compared to Tg Ctrl in the Ent (p = 0.0234). In Tg females, CR induced an opposite effect in the Ent, increasing amyloid burden when compared to female Tg Ctrl (p = 0.0138) and Tg Rsv rats (p = 0.0117).

**Figure 7:**
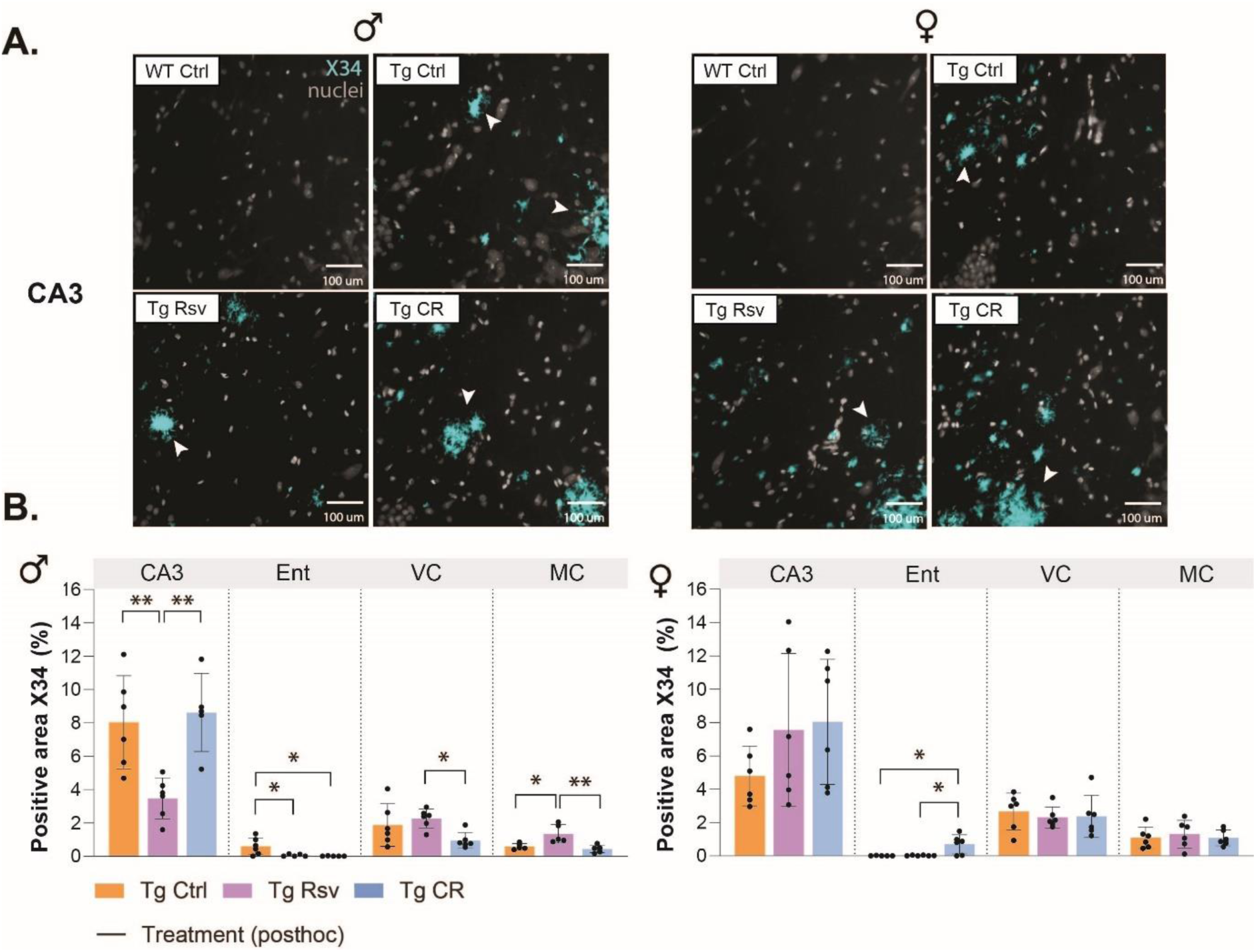
Aβ pathology in male and female, Ctrl, Rsv and CR TgF344-AD rats. A. Representative histological images of Aβ-plaques (light-blue, white arrowheads) in the CA3 for male and female WT Ctrl and Ctrl, Rsv, and CR TgF344-AD rats. Nuclei are counterstained in gray. B. Graphs represent the percentage positive area Aβ, for 4/11 brain regions per sex. Group means ± SD are presented together with the individual subject data points (black dots). Significant treatment effects (one-way ANOVA, post-hoc Tukey HSD p<0.05) are annotated by the black lines. Asterisks indicate the levels of statistical significance: *p<0.05 **p<0.01.

### Long-term dietary intervention with CR or Rsv supplementation has limited effects on neuroinflammation in TgF344-AD rats and WT littermates

Both Rsv and CR have been shown to exert anti-inflammatory effects. To investigate if neuroinflammation was altered due to long-term dietary interventions, immunohistochemistry was performed using IBA-1 (Figure 8A) and GFAP (Figure 9A) for microglia and astrocytes respectively in 11 different brain regions (Supplementary Table 2).

**Figure 8:**
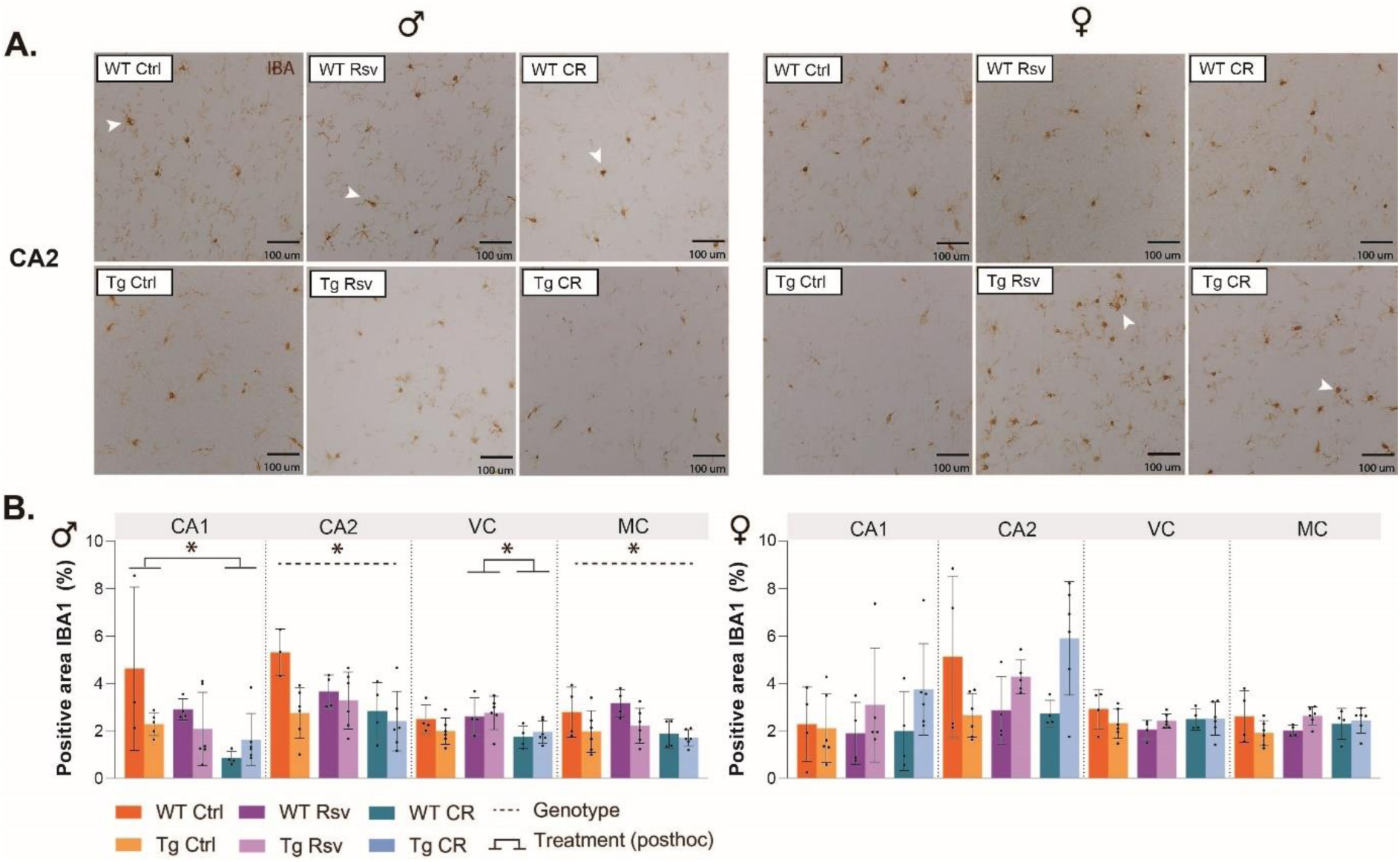
IBA-1-stained microglia in male and female, Ctrl, Rsv and CR TgF344-AD rats. A. Representative histological images show microglia (brown, white arrowheads) in the CA2 for male and female Ctrl, Rsv, and CR WT and TgF344-AD rats. B. Graphs represent the percentage positive area IBA-1, for 4/11 brain regions per sex. Group means ± SD are presented together with the individual subject data points (black dots). Significant genotype effects (two-way ANOVA, p < 0.05) are annotated by the black dotted line. Significant treatment effects (post-hoc Tukey HSD p<0.05) are annotated by the black brackets. Asterisks indicate the levels of statistical significance: *p<0.05.

**Figure 9:**
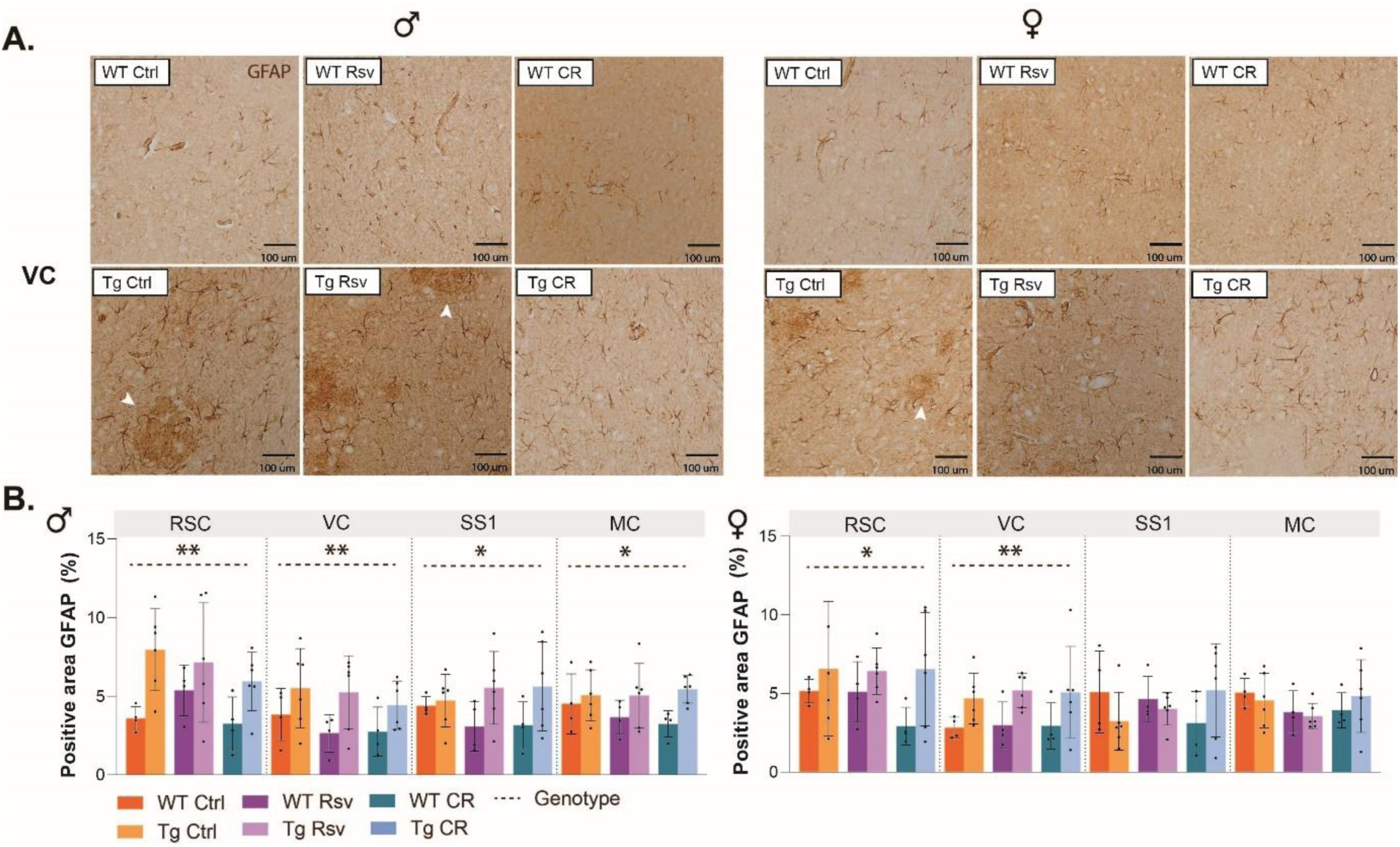
GFAP stained astrocytes in male and female, Ctrl, Rsv and CR TgF344-AD rats. A. Representative histological images show astrocytes (brown, white arrowheads) in the VC for male and female Ctrl, Rsv, and CR WT and TgF344-AD rats. B. Graphs represent the percentage positive area GFAP, for 4/11 brain regions per sex. Group means ± SD are presented together with the individual subject data points (black dots). Significant genotype effects (two-way ANOVA, p < 0.05) are annotated by the black dotted line. Asterisks indicate the levels of statistical significance: *p<0.05 **p<0.01.

Statistical analysis (two-way ANOVA, p < 0.05) of IBA-1 revealed significant treatment effects in males for 2 out of 11 regions (Figure 8B, Supplementary Figure 7). Post-hoc analysis shows lower IBA-1 signal for CR male rats when compared to Ctrl in the CA1 (p = 0.0383), and when compared to Rsv in the VC (p = 0.0135), both irrespective of genotype. Additionally, significant genotypical differences were observed in 2 out of 11 regions, where male WT rats had higher IBA-1 signal compared to Tg in the CA2 (p = 0.0317) and MC (p = 0.0237). In female rats, the IBA-1 signal demonstrated no significant main or interaction effects in any region.

Statistical analysis (two-way ANOVA, p < 0.05) of GFAP revealed significant genotype effects in males for 4 out of 11 regions, and 2 out of 11 for females (Figure 9B, Supplementary Figure 8). In males, we observed higher GFAP signal in Tg rats compared to WT in the RSC (p= 0.0029), VC (p = 0.0088), SS1 (p = 0.0257) and MC (p = 0.0204). For females, similar higher GFAP signal in Tg compared to WT was observed in the RSC (p = 0.0469) and VC (p = 0.0033).

## 4. Discussion

This study investigated the effects of short- and long-term dietary intervention using CR or Rsv supplementation on brain function, cognition and neuropathological markers in male and female TgF344-AD rats and WT littermates. At both ages, CR induces a decrease of rs-FC in males and females. Long-term CR yielded modest cognitive improvements in male Tg rats, as evidenced by enhanced performance in the MWM, but showed limited cognitive benefits in females. Modest effects were observed as a result of long-term CR and Rsv supplementation on amyloid pathology and microgliosis. However, differences in microgliosis and astrogliosis were primarily driven by genotype, with persistent distinctions between WT and Tg rats regardless of treatment.

In our work, we observed male-specific hyperconnectivity, irrespective of treatment, in Tg rats compared to WT at 4M of age, preceding hypoconnectivity observed at 11M of age, while in females at both 4- and 11M of age hypoconnectivity was observed in Tg rats compared to WT. While our results are consistent with the well-documented hypoconnectivity observed at the plaque stages in both patients with MCI/AD (43–45), and rodent models of AD (19–22), they also reflect the inconsistency between observed hypo- and hyperconnectivity reported in the literature at earlier disease stages in patients (46–50) and rodent models of AD. Altogether, our results contribute to a growing body of evidence suggesting a biphasic trajectory of FC changes in early AD.

It is commonly accepted that lifestyle interventions such as a reduction in energy (caloric) intake, or dietary supplementation with Rsv, could contribute to the delay in onset of age-related diseases, including AD. However, only few studies have investigated the relationship between short-term CR, Rsv supplementation and brain function. In our work, short-term CR in 4M old Tg female rats reduced FC compared to Ctlr and Rsv rats in ROI-pairs within and between the Hipp RSN, during a phase when weight loss was still prominent (Supplementary Figures 1B, D). In our previous work in healthy young adult male and female rats, we observed a female-specific decrease of FC as a result of short-term CR and Rsv supplementation within- and between Hipp and SubC RSNs (39). The study of Prehn et al., investigating the effects of intentional weight loss using short-term (3 months) CR in obese postmenopausal women demonstrated decreased hippocampal connectivity directly after the weight loss phase of CR, which did not persist through the weight maintenance phase, one month after 3 month long CR (36). Supporting this pattern, another study found decreased FC following a 6-month hypocaloric Mediterranean diet in humans (51), in line with our current findings. While we observed a genotype-specific decrease of rs-FC in Tg rats after short-term CR, Rsv supplementation did not recreate similar effects. Furthermore in contradiction with our findings, some human studies in healthy older adults (37) and patients with MCI (40) reported increased FC between the hippocampus and cortical regions after short-term (6 months) Rsv supplementation. Altogether, while short-term CR reduces hippocampal rs-FC, the absence of similar effects with Rsv challenges its role as a true CR-mimetic in the context of brain function.

While short-term changes in FC have been observed following CR and Rsv supplementation, evidence of long-term effects are severely lacking. In our work, we observed a treatment-specific genotype difference after CR, where male Tg CR rats displayed lower rs-FC compared to WT CR rats. Moreover, we observed a rs-FC decrease after long-term CR as compared to Ctrl in Tg male rats. Here, long-term Rsv produced minimal effects on FC, in contrast to the observed CR-induced changes. As mentioned earlier, hypoconnectivity is well documented at the plaque stage, correlated to cognitive impairment (52, 53). In this regard, the decrease in rs-FC observed after long-term CR in Tg male rats might prove detrimental in contrast to the expected and hypothesized beneficial outcomes of dietary intervention with CR. To our knowledge, no study has assessed the effects of long-term CR or Rsv supplementation on brain function using rsfMRI in healthy individuals, patients nor rodent models of AD. The scarcity of studies makes it difficult to determine the potential long-term benefits, efficacy and mechanisms of CR and Rsv supplementation, emphasizing the need for longitudinal research to assess their therapeutic potential in aging and neurodegeneration.

Spatial disorientation is a common symptom of AD, often linked to hippocampal dysfunction. The MWM is typically used to study hippocampal-dependent spatial learning and memory in rodents. In our study, Tg CR males showed reduced travel distances, suggesting slightly improved spatial memory, though not enough to confirm cognitive benefits. Moreover, we did not observe any potential beneficial effects for the supposed CR-mimetic Rsv. Additionally, we observed in both male and female rats significant longer distances travelled and escape latencies for Tg rats compared to WT. Currently, only a few studies have investigated spatial memory impairment in the TgF344-AD rat model. During pre- and early-plaque stages (4-6M), male and female WT and Tg rats show little to no difference in MWM performance (54). However, in studies where sexes were combined, Tg rats exhibit increased latency in reversal learning at 6M (55) and 24M (54). At 11M of age, spatial deficits become pronounced, including reduced path linearity, longer swim paths, and escape latencies in Tg rats compared to WT (55), in line with our own findings. Studied effects of CR on MWM performance are limited, with research performed in aged rodents reporting no beneficial effects of CR or Rsv supplementation on cognitive improvement (56). Some studies however found improved performance of CR in F344 rats (57) and AD mice (58, 59), as they travelled less distance within the maze and had lower escape latencies compared to their ad-libitum fed littermates, similar to our observations. Findings on the efficacy of Rsv remain mixed, with some studies report improved spatial memory (60–63) and others reporting impaired learning (60–62), while in our work, Rsv showed negligible effects on spatial memory. Differences in outcomes may result from administration methods (i.p. versus oral). Studies have shown that the oral bioavailability of resveratrol is low (64). Still, due to its lipophilic nature, Rsv can cross the blood–brain barrier (65), as (very low) levels have been detected in neural tissue (66–68), supporting the idea that even low concentrations may underlie the reported neuroprotective effects of Rsv. Given the inconsistent results and low bioavailability of Rsv, more rigorous, long-term studies with larger sample sizes, direct measurement of Rsv brain levels, and controlled administration methods are needed to better determine the effectiveness of these interventions.

Our findings show that long-term CR or Rsv supplementation modulates amyloid plaque load in TgF344-AD rats in a region-specific manner. Research in rodent models of AD has been able to show attenuation of amyloid plaque accumulation as a result of CR (25, 27–29) or Rsv supplementation (35, 66, 69), primarily within hippocampal regions. In line with these findings, we observed an attenuation of amyloid plaque load in Rsv supplemented Tg rats when compared to Tg Ctrl and Tg CR rats in the CA3 and Ent. CR has been widely speculated to exert its anti-amyloidogenic effects through enhancement of autophagy, a process known to degrade aggregated proteins including Aβ (70), while Rsv has been speculated to exert its effects through direct competition with Aβ plaque binding sites (71), enhancing Aβ peptide cleavage (34) and reducing fibril length (33). As mechanisms vastly differ between CR and its mimetic, Rsv, there is a potential underlying hypothesis explaining the differential effects of both interventions that we observe within our data. Moreover, in the VC and MC in males, we observe an increased amyloid burden upon Rsv supplementation, in contrast with the observed attenuation of amyloid plaque load in the CA3 and Ent. Despite its reported benefits on Aβ, research has also shown that Rsv is unable to prevent Aβ oligomerization (72). Additionally, Rsv inhibits ubiquitin-dependent protein degradation (73), blocking critical regulators of Aβ clearance, potentially resulting in increased amyloid burden. Taken together, while CR and Rsv have been proposed as potential strategies to reduce Aβ pathology, our results suggest their effects are limited and may vary by brain region, with some areas even showing increased amyloid accumulation as a result of dietary intervention, underscoring the difficulty of altering key disease processes with these long-term dietary approaches.

While Aβ accumulation has long been a central focus in AD research, increasing evidence highlights the contributory role of neuroinflammatory processes in both the onset and progression of the disease, with chronic activation of microglia and astrocytes contributing to neuronal dysfunction, synaptic loss, and exacerbation of amyloid and tau pathology (74, 75). In male CR rats, we observe less microglia in the CA1 compared to Ctrl, however, in regions such as the CA2 and MC CR does not alter microglial levels. Moreover, Rsv supplementation fails to alter microglia levels. CR and Rsv are both heavily implied in delaying the onset and attenuation of progression of AD through their anti-inflammatory effects (76, 77). CR has been shown to reduce the number of hypothalamic microglia in aging (78) and hippocampal microglia in AD APP_swe_/PS1_delta9_ mice (29), while failing to exert changes in microglial levels in the corpus callosum of aged monkey brain (79) and in the hippocampus of AD APP*_swe/ind_* mice (25). These mixed findings suggest that the anti-inflammatory effects of CR, particularly on microglia, may be region-, age-, and model-specific. Similarly, the evidence regarding the effects of Rsv on microglia is complex and sometimes contradictory. Rsv supplementation was able to attenuate microglial activation in animal models of neurodegenerative pathologies (80, 81), including an amyloid β25-35 induced AD rat model (63). Studies have shown that Rsv has an impact on microglial transcriptomes (82), speculated through regulation of inflammatory and oxidative regulators such as IL-1β and IL-6, inducible nitric oxide synthase (iNOS) (82) and SIRT1 (83). In a study by Porquet et al., long-term Rsv supplementation (10 months) did not alter inflammatory and oxidative mechanisms in a mouse model of familial AD (35). However, the authors do speculate that there is a possibility that Rsv exerts earlier antioxidant effects at younger ages, acting more as a preventive than as a curative agent. Within the same work, Porquet et al report a decreased SIRT1 activation after Rsv supplementation. SIRT1 deficiency in microglia has been implicated in cognitive decline and neurodegeneration (84). A recent meta-analysis by Mansouri et al. (85) has concluded that despite the believed influence of Rsv on SIRT1, supplementation has its limited effects on SIRT1. However, their dose-response analysis suggests that the effect of Rsv is dependent on the dosage regimen, which could potentially underlie these reported differences. Rsv is known to exert a hormesis dose-dependent biphasic effect (86, 87). Therefore, the absence of microglial modulation by Rsv observed in our study may be attributed to suboptimal dosing. This highlights the need for future studies to systematically evaluate dosage regimens to better understand and optimize the therapeutic potential of Rsv in neurodegenerative disease models.

In our work, CR and Rsv supplementation do not appear to alter astrocyte levels. CR has been reported to be able to remodel astrocytes (88, 89), contributing to enhanced synaptic plasticity. However, short-term CR failed to alter astrocyte levels in AD mice, unless proximity to amyloid plaques was taken into consideration (25), what despite the difference in treatment duration, is somewhat in line with our own findings. Short-term Rsv supplementation has been shown to reduced astrocyte levels in an amyloid-β25-34 induced rat model (63). Similar to the effects of CR on astrocytes, Rsv seems to impact more specifically astrocyte functional remodeling (90) and inflammation-related gene expression (91). Overall, these findings suggest that both CR and Rsv primarily influence astrocyte function, while Rsv additionally has the potential to affect astrocyte levels. Future work should focus on spatial and functional characterization of astrocytes to better understand how CR and Rsv influence their role in AD pathology.

Based on well-established sex-differences in context of AD (92–94), our work deliberately characterized males and females separately, aiming to provide more insight into sex-specific AD pathology of the TgF344-AD rat model and the potential sex-specific impact of CR and Rsv (39) on disease progression. Risk factors known to increase prevalence and progression of AD in women specifically include (but are not limited to) deviations in brain structure and function, molecular biomarkers and inflammation. Men have a larger brain volume on average, and thus are less sensitive to molecular pathology in AD and suffer less or slower structural loss as compared to women (95, 96), which are known to have higher amyloid burden (97) and levels of tau tangle density (98). Moreover, males have stronger interhemispheric FC within the hippocampus compared to women. Women usually have stronger immune responses to stimuli than men, with involvement of different pathways and immune cells (99). For example, glial cell-mediated neuro-immune responses have been found to show sex-specific dysregulation, implicated in AD (100). In 5xFAD and 3xTg-AD mouse models of AD, most of these findings are reiterated, with female Tg mice showing higher amyloid accumulation, NFTs, neuroinflammation and spatial cognitive deficits compared to male mice (101–103).

This research is subject to some limitations. First, this study is limited by its sample size. Power calculations showed sufficient power upon pooling of both males and females. However, as we proved sex-specific effects of CR or Rsv supplementation in our previous work (39), the decision was made to consider the sexes separately. Second, both CR and Rsv are both known to influence the vasculature (104–106). The BOLD signal, used in this study as an indirect measure of neuronal activity, is heavily dependent on vascular parameters such as CBF, cerebral blood volume (CBV) and the rate of oxygen consumption in response to changes in neuronal activity (107–109). Considering the potential systemic vascular effects of CR and Rsv, it is possible they contribute to changes in the BOLD signal, which should be considered when interpreting BOLD outcomes in comparison to Ctrl. Third, while measures like escape latency and path length are useful for assessing cognition and treatment effects, they often overlook animals’ swimming behavior and search strategies. Animals use various strategies to locate the platform, prompting the development of more refined analysis methods (110, 111). In mouse models of AD, such analysis has revealed cognitive differences missed upon utilization of basic measures, with Tg mice using more random, non-spatial strategies than WT (112, 113). These findings highlight the value of detailed behavioral assessments and suggest that applying similar methods in our study could uncover subtle cognitive changes and treatment effects in the TgF344-AD rat model.

In conclusion, our work demonstrated the effects of short- and long-term CR and Rsv supplementation on brain function, cognition, and AD-related pathology in male and female TgF344-AD rats and WT littermates. While short- and long-term Rsv showed minimal impact across most parameters, CR induced a reduction in rs-FC as compared to Ctrl rats for many connections mainly in female rats upon short -term CR, and male rats upon long-term CR. Additionally, both short- and long-term CR reduced rs-FC for specific ROI-pairs in Tg rats only. Long-term CR induced modest cognitive improvements in spatial memory. Amyloid pathology and neuroinflammation were largely unaffected as a result of long-term dietary intervention with CR or Rsv supplementation, though region-specific reductions in microglial activation were observed in male CR rats specifically. These findings suggest that CR may modulate early and late AD-related pathology, while Rsv is unable to recapitulate these effects. Our findings underscore the importance of sex as a biological variable and highlight the need for further research into the mechanisms of CR or Rsv supplementation as intervention to delay or attenuate AD progression.

## Author contributions (CRediT)

JR: conceptualization, data curation, formal analysis, investigation, project administration, validation, visualization, writing-original draft, writing – review& editing. MB: conceptualization, formal analysis, project administration, software, supervision, validation, visualization, writing-review & editing. MVV: data curation, formal analysis, investigation, visualization, validation. EC: conceptualization, data curation, investigation, software, validation. TV: formal analysis, software, supervision, validation, investigation, writing – review & editing. LK: conceptualization, investigation. IvS: methodology, resources, software. JA: methodology, resources, software. DvD: methodology, resources, software, supervision, writing – review & editing. DB: supervision, visualization, writing – review & editing. MA: methodology, resources, software, supervision, validation, visualization, writing – review & editing. MV: conceptualization, methodology, project administration, resources, supervision, validation, writing – review & editing.

## Funding

This study was supported by the Fund of Scientific Research Flanders (FWO-G045420N) and Stichting Alzheimer Onderzoek (SAO-FRA 2020/027). The computational resources and services used in this work were provided by the HPC core facility CalcUA of the University of Antwerp, the VSC (Flemish Supercomputer Center), funded by the Hercules Foundation and the Flemish Government department EWI. Funding for heavy scientific equipment was provided by the Flemish Impulse funding under grant agreement number 42/FA010100/1230.

## Acknowledgements

We would like to acknowledge Dr. Georgios A. Keliris for his initial contributions to the project proposal, which lead to the SAO-granted funding.

## Notes

### Competing Interest Statement

The authors have declared no competing interest.

