## Supplementary material for "Long-term dietary interventions fail to mitigate functional connectivity loss and cognitive decline in the TgF344-AD rat model of Alzheimer’s Disease"

### SUPPLEMENTARY INFORMATION

**Supplementary Table 1.** Brain regions and resting-state network abbreviations

| Resting-state network | Region | Abbreviation |
|---|---|---|
| **Default mode-like network (DMLN)** | Cingulate cortex left | Cg L |
| | Cingulate cortex right | Cg R |
| | Retrosplenial cortex left | RSC L |
| | Retrosplenial cortex right | RSC R |
| | Prelimbic cortex left | Limbic L |
| | Prelimbic cortex right | Limbic R |
| | Temporal Association left | TeA L |
| | Temporal Association right | TeA R |
| | Infralimbic cortex left | IL L |
| | Infralimbic cortex right | IL R |
| **Hippocampal network (Hipp)** | Hippocampal field CA1 left | CA1 L |
| | Hippocampal field CA1 right | CA1 R |
| | Hippocampal field CA2 left | CA2 L |
| | Hippocampal field CA2 right | CA2 R |
| | Hippocampal field CA3 left | CA3 L |
| | Hippocampal field CA3 right | CA3 R |
| | Dentate gyrus left | DG L |
| | Dentate gyrus right | DG R |
| | Entorhinal cortex left | Ent L |
| | Entorhinal cortex right | Ent R |
| **Sensory network (Sens)** | Visual cortex left | VC L |
| | Visual cortex right | VC R |
| | Auditory cortex left | Aud L |
| | Auditory cortex right | Aud R |
| | Piriform cortex left | Pir L |
| | Piriform cortex right | Pir R |
| **Lateral cortical network (LCN)** | Primary somatosensory cortex left | S1 L |
| | Primary somatosensory cortex right | S1 R |
| | Secondary somatosensory cortex left | S2 L |
| | Secondary somatosensory cortex right | S2 R |
| | Motor cortex left | MC L |
| | Motor cortex right | MC R |
| | Frontal association cortex left | FrA L |
| | Frontal association cortex right | FrA R |
| | Insular cortex left | Ins L |
| | Insular cortex right | Ins R |

| | | |
|---|---|---|
| **Subcortical network (SubC)** | Caudate putamen left<br>Caudate putamen right<br>Nucleus Accumbens left<br>Nucleus Accumbens right<br>Basal forebrain<br>Thalamus left<br>Thalamus right<br>Superior colliculus left<br>Superior colliculus right<br>Hypothalamus left<br>Hypothalamus right | CPu L<br>CPu R<br>NAcc L<br>NAcc R<br>BFB<br>Thal L<br>Thal R<br>SC L<br>MS R<br>Hyp L<br>Hyp R |

**Supplementary Table 2.** Brain regions selected for histological analysis and their abbreviations

| Region | Abbreviation |
|---|---|
| Cingulate cortex | Cg |
| Retrosplenial cortex | RSC |
| Hippocampal field CA1 | CA1 |
| Hippocampal field CA2 | CA2 |
| Hippocampal field CA3 | CA3 |
| Dentate gyrus | DG |
| Entorhinal cortex | Ent |
| Visual cortex | VC |
| Primary somatosensory cortex | S1 |
| Motor cortex | MC |
| Caudate putamen | CPu |

**Supplementary Table 3.** Statistical outcomes (one-way ANOVA, treatment) of X34 for each measure per sex.

| | **One-way ANOVA** | | | | | | | |
|---|---|---|---|---|---|---|---|---|
| | **Males** | | | | **Females** | | | |
| **Regions** | **DF** | **Sum of squares** | **F ratio** | **P-value** | **DF** | **Sum of squares** | **F ratio** | **P-value** |
| **CA1** | *2* | *3.786* | *1.083* | *0.3674* | *2* | *10.76* | *0.7992* | *0.4692* |
| **CA2** | *2* | *56.15* | *2.545* | *0.1141* | *2* | *8.430* | *0.6751* | *0.5261* |
| **CA3** | *2* | *91.56* | *9.386* | *0.0026** | *2* | *37.12* | *1.451* | *0.2654* |
| **DG** | *2* | *13.82* | *1.544* | *0.2456* | *2* | *13.59* | *1.337* | *0.2943* |
| **Ent** | *2* | *1.202* | *5.973* | *0.0145** | *2* | *1.792* | *7.548* | *0.006** |
| **Cg** | *2* | *0.0993* | *1.926* | *0.1824* | *2* | *0.1364* | *1.621* | *0.2305* |
| **RSC** | *2* | *2.360* | *1.147* | *0.2731* | *2* | *2.729* | *0.4961* | *0.6186* |
| **V** | *2* | *5.522* | *3.798* | *0.0463** | *2* | *0.4417* | *0.2066* | *0.8156* |
| **S1** | *2* | *0.3512* | *0.6997* | *0.5145* | *2* | *3.072* | *3.31* | *0.0665* |
| **M** | *2* | *2.384* | *9.112* | *0.0039** | *2* | *0.2115* | *0.2398* | *0.7897* |
| **CPu** | *2* | *<0.0001* | *2.050* | *0.1715* | *2* | *<0.0001* | *0.7561* | *0.4866* |

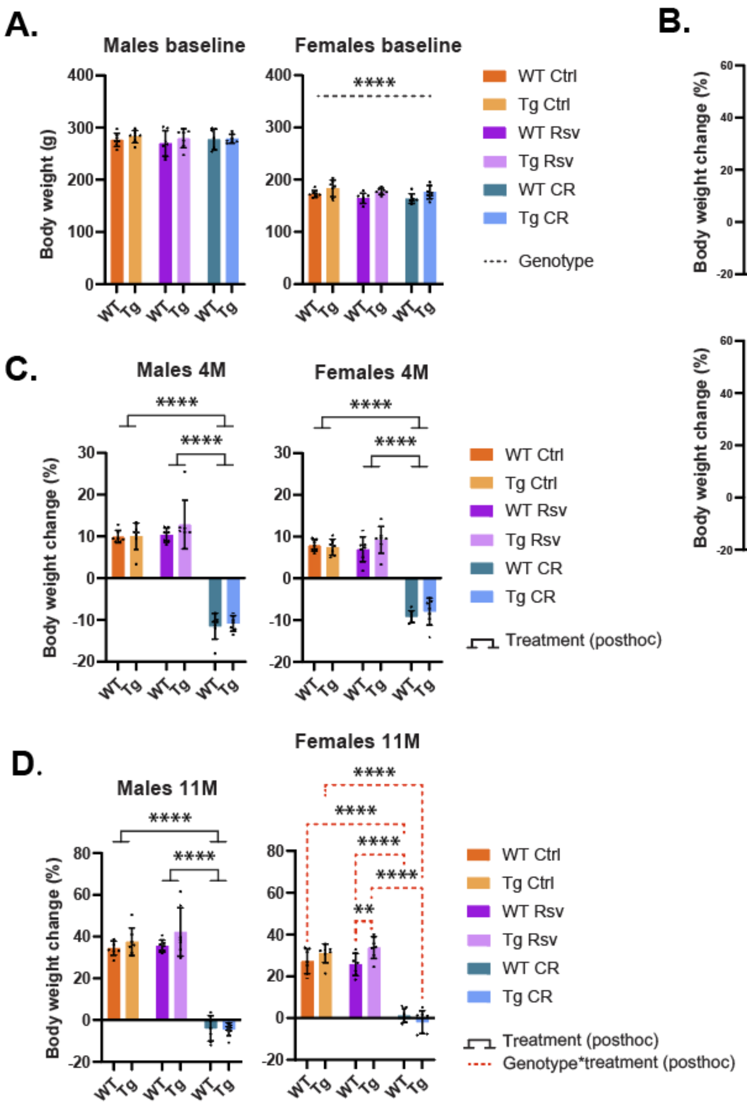


**Supplementary figure 1:** *Effects of dietary intervention on body weight (BW). (A) Mean ± SD BW in grams (g) at baseline, prior to the start of the dietary interventions (week 0) for both males and females. The colors of the bars are indicative of the group each subject was assigned to (control (Ctrl), resveratrol (Rsv), caloric restricted (CR). Dots represent individual subject data points. Asterisks indicate the levels of statistical significance: ****p<0.0001. (B) Percentage change in BW (mean ± SD) with respect to baseline, over the course of the experiment for the Ctrl, Rsv and CR group – both males and females. (C, D) Mean ± SD percentage change in body weight for both male and females at 4M of age (C) and 11M of age (D). Asterisks indicate the levels of statistical significance: **p<0.01, ****p<0.0001.*

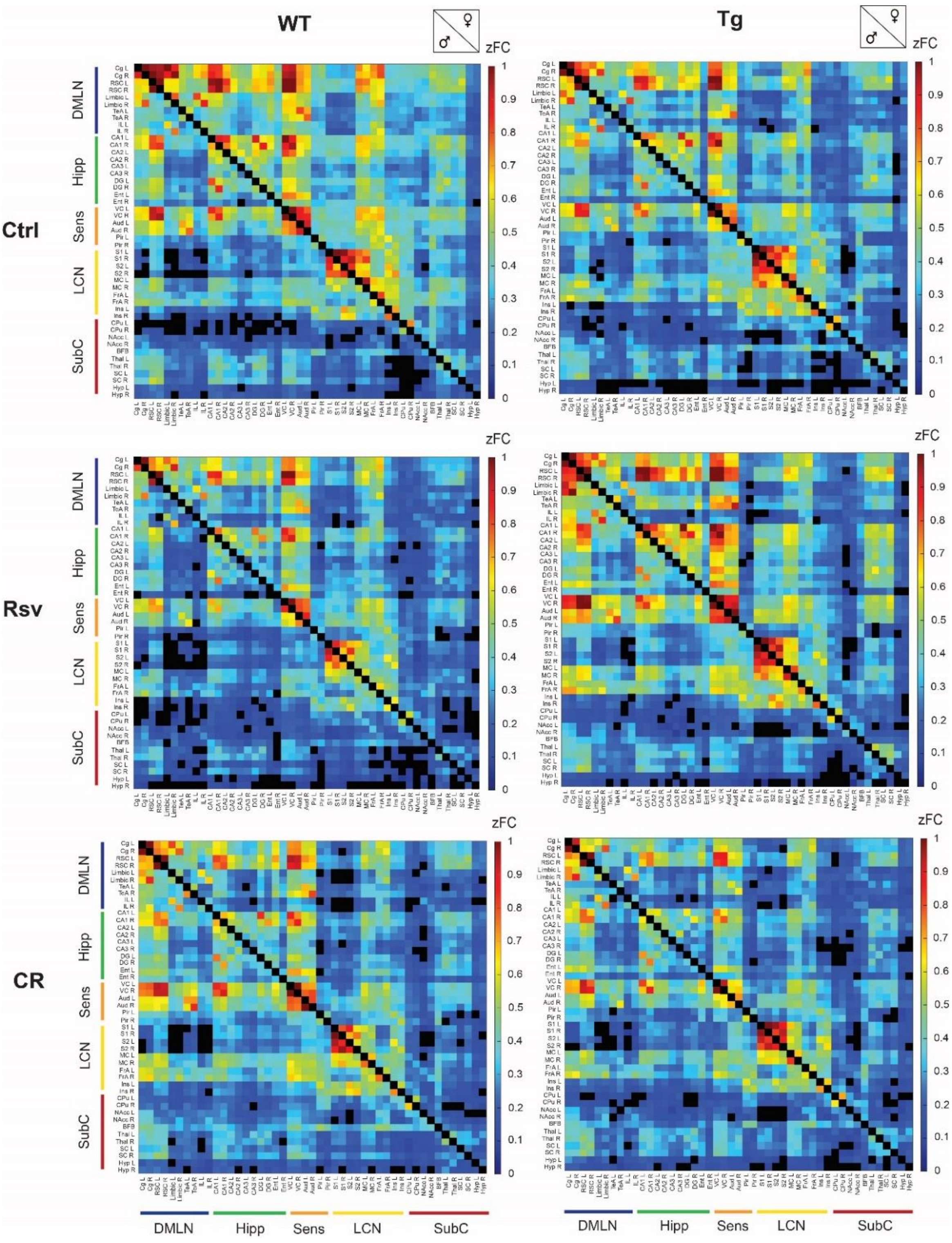


**Supplementary figure 2:** *FC for Ctrl, Rsv supplemented and CR WT and TgF344-AD rats. ROI-based FC matrices displaying the mean zFC after short-term dietary intervention between ROI pairs for WT (lower diagonal) and Tg (upper diagonal) male and female rats per treatment group (Ctrl = control, CR = caloric restriction, Rsv = Resveratrol, zFC = Fisher's z-transformed functional connectivity). The 47 ROIs with left (L) and right (R) hemispheric are grouped together based on their associated RSNs (Supplementary Table 1). Colors represent the mean FC between different significantly connected ROIs. Non-significant connections (p≥ 0.05, one-sample t-test, FDR corrected per group) are blacked out.*

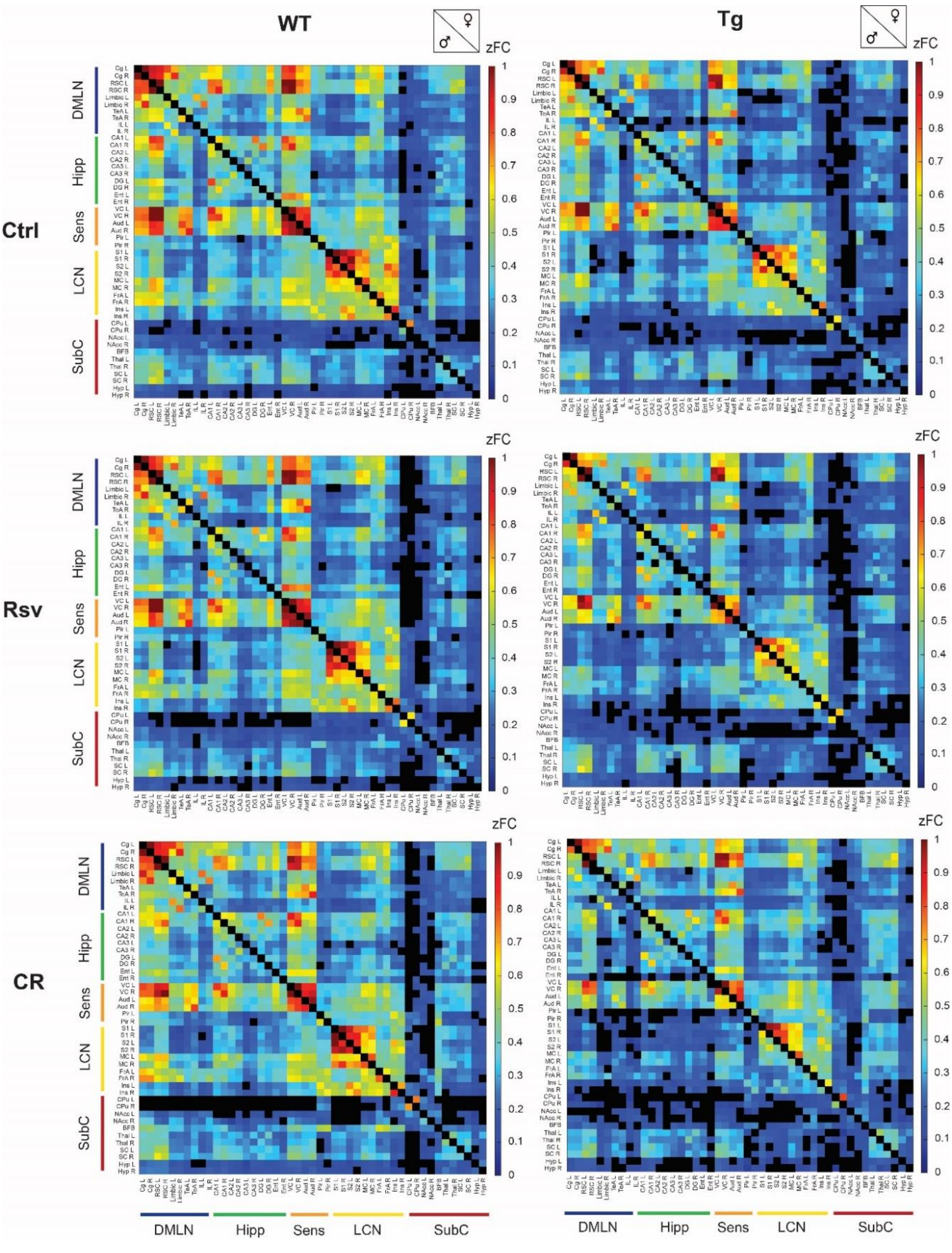


**Supplementary figure 3:** *zFC for Ctrl, Rsv supplemented and CR WT and TgF344-AD rats. ROI-based FC matrices displaying the mean zFC after long-term dietary intervention between ROI pairs for WT (lower diagonal) and Tg (upper diagonal) male and female rats per treatment group (Ctrl = control, CR = caloric restriction, Rsv = Resveratrol, zFC = Fisher's z-transformed functional connectivity). The 47 ROIs with left (L) and right (R) hemispheric are grouped together based on their associated RSNs (Supplementary Table 1). Colors represent the mean zFC between different significantly connected ROIs. Non-significant connections (p≥ 0.05, one-sample t-test, FDR corrected per group) are blacked out.*

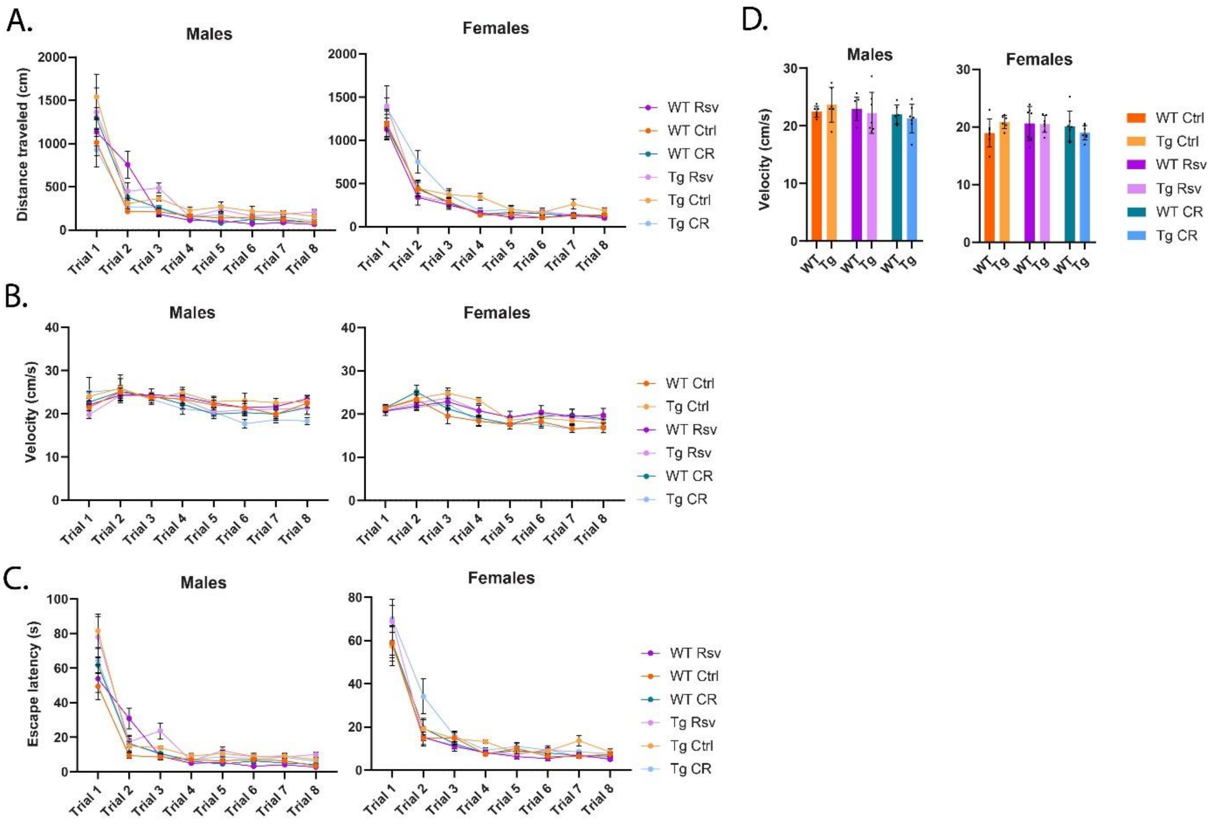


**Supplementary figure 4:** *MWM learning trials for both male and female Ctrl, Rsv supplemented and CR WT and TgF344-AD rats. Total path lengths (A), swim speed (B), escape latency (C) during MWM training trials and respective sums of average swim speeds (D) averaged across all trails. Group means ± SD are presented together with the individual subject data points (dots, bar plots).*

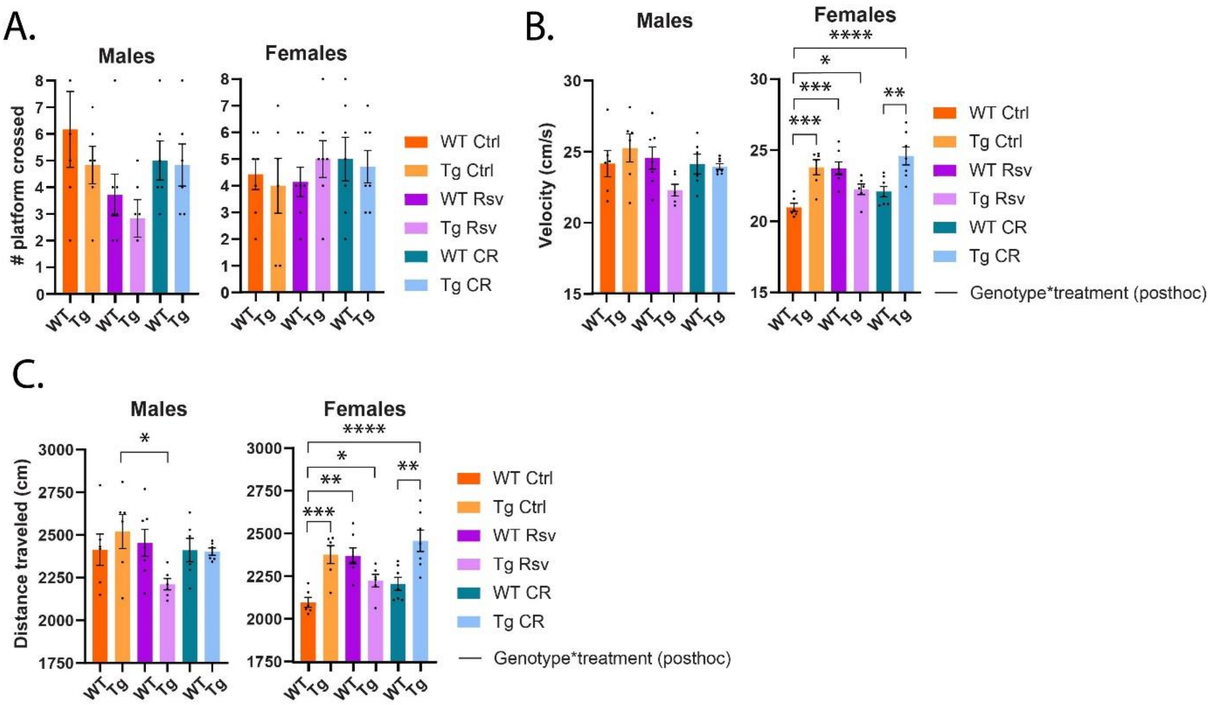

**Supplementary figure 5:** *MWM probe trial outcomes for both male and female Ctrl, Rsv supplemented and CR WT and TgF344-AD rats. The number of times the prior platform location was crossed (B), swim speeds (C) and total distance travelled (D) during the MWM probe trails. Group means ± SD are presented together with the individual subject data points (dots). Significant treatment effects (post-hoc Tukey HSD p<0.05) are annotated by the black brackets. Significant interaction effects of treatment*genotype (FDR corrected, Benjamini-Hochberg procedure, p<0.05) are noted by the black lines. Asterisks indicate the levels of statistical significance: *p<0.05 **p<0.01, ***p<0.001, ***p<0.0001.*

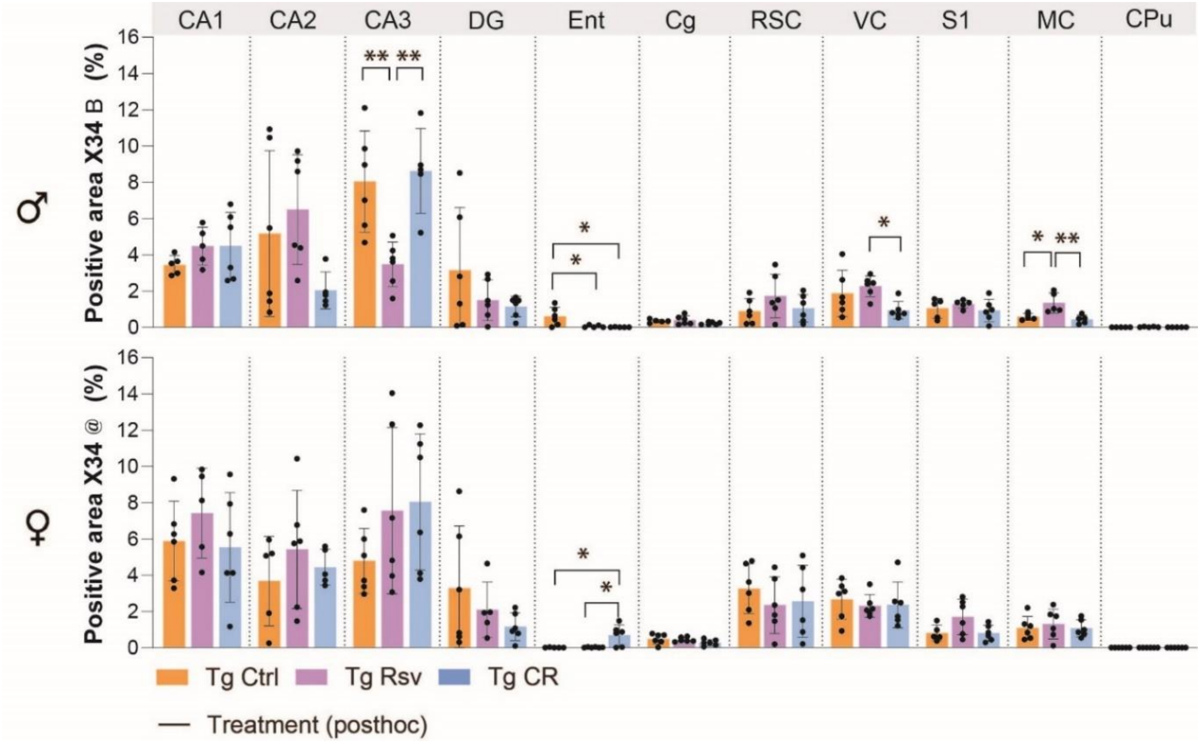


**Supplementary figure 6:** *Percentage positive area X34 for both male and female, Ctrl, Rsv supplemented and CR TgF344-AD rats. Group means ± SD are presented together with the individual subject data points (dots). Asterisks indicate the levels of statistical significance: *p<0.05 **p<0.01.*

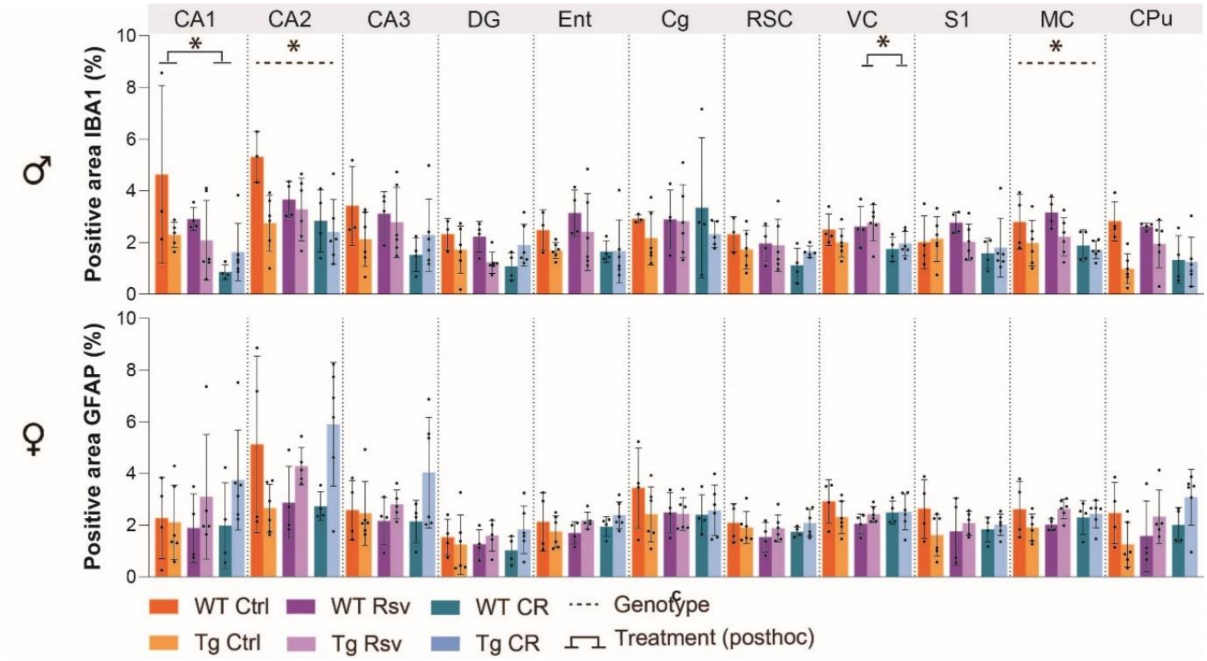


**Supplementary figure 7:** *Percentage positive area IBA1 for both male and female, Ctrl, Rsv supplemented and CR TgF344-AD rats. Group means ± SD are presented together with the individual subject data points (dots). Asterisks indicate the levels of statistical significance: *p<0.05.*

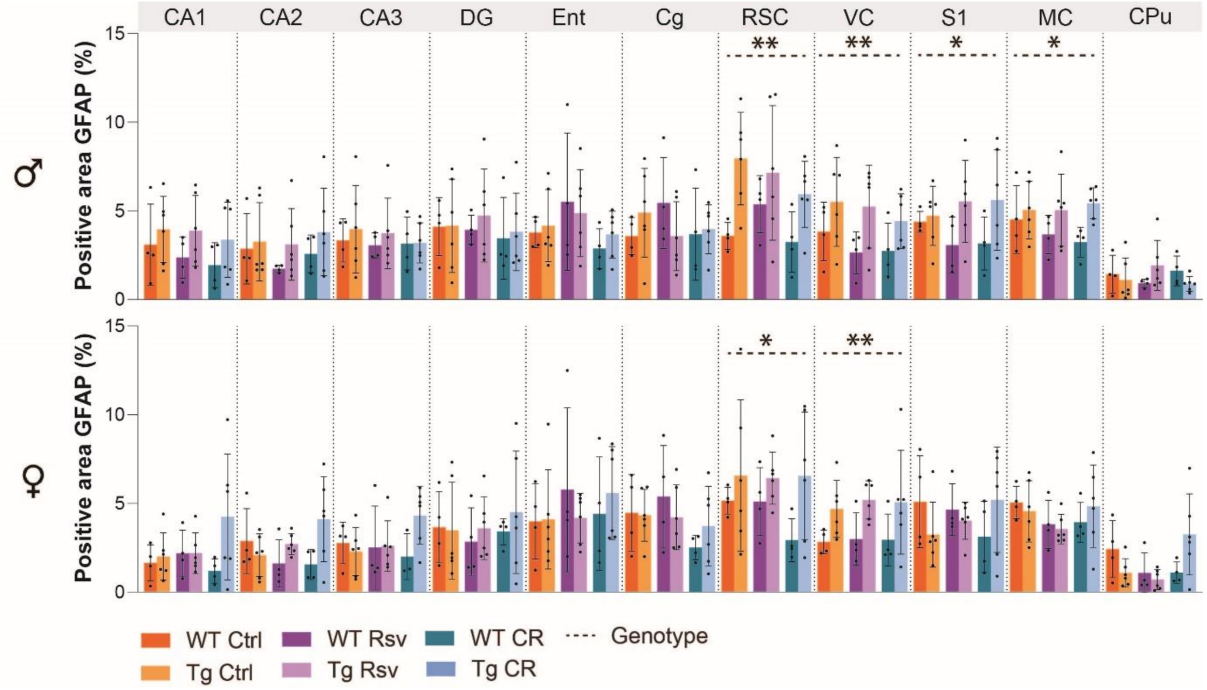


**Supplementary figure 8:** *Percentage positive area GFAP for both male and female, Ctrl, Rsv supplemented and CR TgF344-AD rats. Group means ± SD are presented together with the individual subject data points (dots). Asterisks indicate the levels of statistical significance: *p<0.05, **p<0.01.*
